# Evolutionary rescue can prevent rate-induced tipping

**DOI:** 10.1101/2020.12.13.422565

**Authors:** Anna Vanselow, Lukas Halekotte, Ulrike Feudel

## Abstract

Today, the transformation of ecosystems proceeds at unprecedented rates. Recent studies suggest that high rates of environmental change can cause *rate-induced tipping*. In ecological models, the associated *rate-induced critical transition* manifests during transient dynamics in which populations drop to dangerously low densities. In this work, we study how *indirect evolutionary rescue* - due to the rapid evolution of a predator’s trait - can save a prey population from the rate-induced collapse. Therefore, we explicitly include the time-dependent dynamics of environmental change and evolutionary adaptation in an eco-evolutionary system. We then examine how fast the evolutionary adaptation needs to be to counteract the response to environmental degradation and express this relationship by means of a critical rate. Based on this critical rate, we conclude that indirect evolutionary rescue is more probable if the predator population possesses a high genetic variation and, simultaneously, the environmental change is slow. Hence, our results strongly emphasize that the maintenance of biodiversity requires a deceleration of the anthropogenic degradation of natural habitats.

## Introduction

Today, populations are faced with unprecedented rates of global climate change, habitat fragmentation and destruction causing an accelerating conversion of their living conditions (Frishkoff et al., 2016; Lee et al., 2017; Oliver et al., 2015; Parmesan, 2006; Selwood et al., 2015; Teyssèdre and Robert, 2014; Trathan et al., 2015; Travis, 2003). In many cases, populations are unable to counter these high rates of change due to i.e. barriers preventing dispersal to more suitable habitats or insufficient time to adapt via phenotypic plasticity or genetic variations (Urban et al., 2016; Williams et al., 2008). Consequently, affected populations can shrink to dangerously low densities or even become extinct (Johnson et al., 2017). Such drastic changes of population densities are called *regime shifts* (Folke et al., 2004; Hastings et al., 2018; Scheffer et al., 2001; Wernberg et al., 2015) or *tipping events* (Ashwin et al., 2012; Siteur et al., 2016; Vanselow et al., 2019; Wieczorek et al., 2011).

In ecology, regime shifts have often been associated with transitions of ecosystems from one stable state to another *alternative stable state* (Dwomoh and Wimberly, 2017; Gårdmark et al., 2015; van de Leemput et al., 2016). So far, based on the existence of alternative stable states, three types of regime shifts or tipping events have been formulated whose course can best be illustrated using potential landscapes (Scheffer et al., 2008) (see fig. 1A-C): (i) *bifurcation-induced tipping* (B-tipping) where slowly changing environmental conditions alter the potential landscape until the formerly stable state (left valley) vanishes and the system state (ball) proceeds to an alternative stable state (right valley) (fig. 1A) (Ashwin et al., 2012; Folke et al., 2004); (ii) *noise-induced tipping* (N-tipping) where the state of the system (ball) is pushed out of the basin of attraction of one stable state (left valley) by fluctuations (noise, fig. 1B) (Ashwin et al., 2012; Horsthemke and Lefever, 1984); and finally (iii) *shock-induced tipping* (S-tipping) where a single large perturbation (shock, extreme event) kicks the system out of its basin of attraction (fig. 1C) (Halekotte and Feudel, 2020).

**Figure 1:**
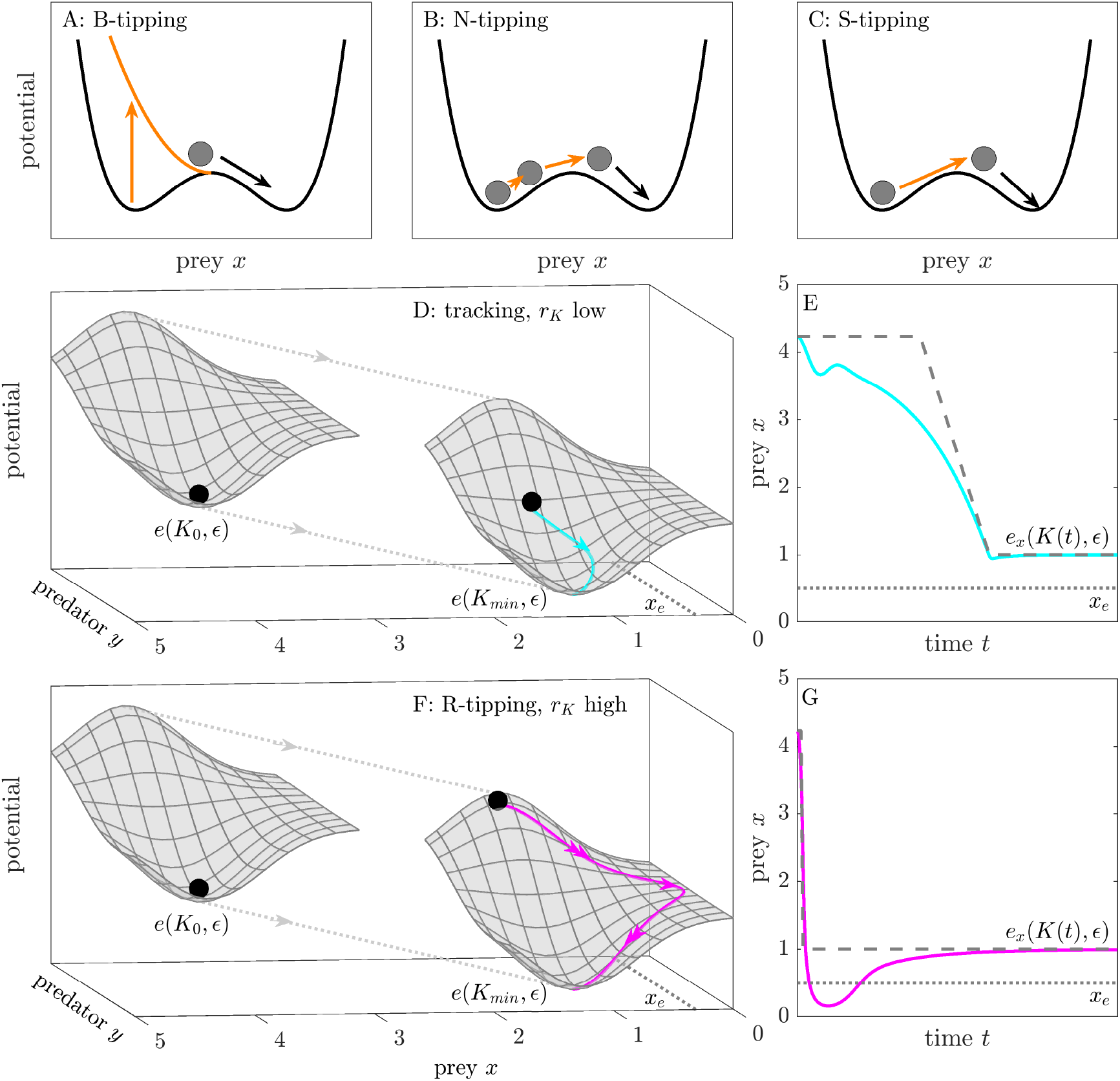
Illustrating B-tipping (A), N-tipping (B), S-tipping (C) using a **sketch** of a two-dimensional potential landscape. (D,F) Demonstrating tracking/R-tipping of the slow-fast predator-prey system (1)–(2) by employing a **sketch** of its three-dimensional potential landscape when *K*(*t*) is fixed at *K*_0_ and *K*_*min*_. (E) Tracking (*r*_*K*_ low): prey density *x* remains above extinction threshold *x*_*e*_ for all time (cyan trajectory). (G) R-tipping (*r*_*K*_ high) causes prey densities to drop below *x*_*e*_ (magenta trajectory).

However, a fourth critical transition is possible which - in contrast to the other three - does not require necessarily the presence of alternative stable states (fig. 1D–G). Instead, it manifests during the transient dynamics – the dynamics before the long-term dynamics are reached (Hastings et al., 2018) – as a system performs a large excursion in state space when external conditions change too fast (Ashwin et al., 2012). During this excursion, it can embrace dangerously, unexpected states as, for instance, a temporarily increase in CO_2_-concentrations in the atmosphere (Wieczorek et al., 2011) or a rapid population collapse (Siteur et al., 2016; Vanselow et al., 2019). For this reason, this tipping mechanism might provide an explanation for regime shifts that occur on comparatively short and therefore ecologically relevant time scales. In reference to the *critical rate* of change which has to be surpassed, this transition is called *rate-induced tipping* (R-tipping).

In order to demonstrate how R-tipping manifests in unexpected - possibly dramatic - transient dynamics, we exemplary study a slow-fast predator(*y*)-prey(*x*) system (1)–(2) that is exposed to environmental change at a given rate. The different time scales attribute to the allometric scaling relationship between different trophic levels (Arditi and Ginzburg, 1989). Here, the prey population evolves faster than its predator which is expressed by the time-scale parameter: 0 < *ϵ* ≪ 1.

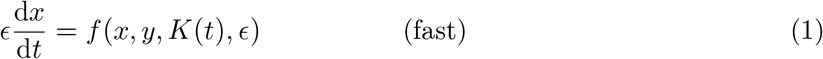

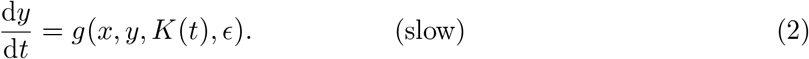

In the following, this predator-prey system is exposed to a gradual environmental change which we model by linearly decreasing the parameter *K* in time at the rate *r*_*K*_ until a prescribed minimum *K*_*min*_ > 0 is reached (see fig. 3C):

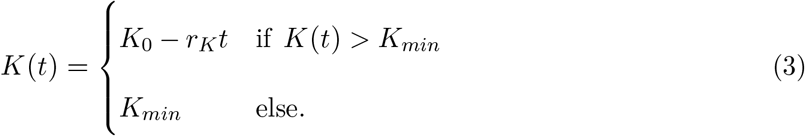

Accordingly, the gradual environmental change is temporary and ends as soon as *K*(*t*) = *K*_*min*_. Furthermore, we assume that for all values of the time-dependent parameter *K*(*t*) ∈ [*K*_0_ *K*_*min*_], the system (1)–(2) possesses an unique stable equilibrium state ***e***(*K*(*t*), *ϵ*) = (*e*_*x*_(*K*(*t*)*ϵ*), *e*_*y*_(*K*(*t*), *ϵ*)) resulting from *f* (*x, y, K*(*t*), *ϵ*) = *g*(*x, y, K*(*t*), *ϵ*) = 0. Due to its dependence on *K*(*t*), ***e***(*K*(*t*), *ϵ*) changes its position in phase space over time. For this reason, ***e***(*K*(*t*), *ϵ*) is called a *moving equilibrium*. Often, such equilibria are also referred to as *quasi-static* which stems from the assumption that they would be adopted if the ecosystem dynamics evolves much faster than the environmental change (Ashwin et al., 2012). Ultimately, as *K*(*t*) = *K*_*min*_ is reached, the system inevitably approaches the sole stable long-term state ***e***(*K*_*min*_, *ϵ*). We choose *K*_*min*_ in such a way that the equilibrium prey densities *e*_*x*_(*K*(*t*), *ϵ*) and *e*_*x*_(*K*_*min*_, *ϵ*) remain above a defined extinction threshold *x*_*e*_ for all times (see fig. 3). Such a threshold can be seen as a ‘rule-of-thumb’ above which it is reasonable to ignore demographic stochasticity (Gomulkiewicz and Holt, 1995) and thus a population whose density remains above *x*_*e*_ is in no danger of becoming extinct. However, an equilibrium density above *x*_*e*_ does not ensure that the system is ‘safe’ as it could have gone through a phase of endangerment during the transient – the dynamics before it settled on ***e***(*K*_*min*_, *ϵ*). This will be demonstrated in the following.

In accordance with B-, N- and S-tipping in figure 1A–C, we employ sketches of the potential landscape of the predator-prey system (1)–(2) to exemplify the differences in its transient dynamics for low (fig. 1D) and high rates of environmental change *r*_*K*_ (fig. 1F). Notice that in both cases start- and end-position of ball and potential are identical. In the beginning *K*(0) = *K*_0_, the ball resides in the minimum of the potential. Hence, the ecosystem is in equilibrium: *x*(0) = *x*_0_ = *e*_*x*_(*K*_0_, *ϵ*) and *y*(0) = *y*_0_ = *e*_*y*_(*K*_0_, *ϵ*). Then, as *K*(*t*) decreases at the rate *r*_*K*_, the system state (ball) follows the moving potential and finally, after *K*(*t*) has reached its minimum *K*_*min*_, settles on ***e***(*K*_*min*_, *ϵ*). However, due to the ‘reaction time’ of the system, the potential slides beneath the ball creating a gap between the current system state (ball) and the moving minimum of the potential ***e***(*K*(*t*), *ϵ*). The size of this gap naturally depends on the speed or rate of environmental change (*r*_*K*_ low/high). As shown by the cyan and magenta trajectory, different gap sizes cause different pathways along which the ball approaches the long-term state *e*(*K*_*min*_, *ϵ*). While the cyan trajectory remains in the vicinity of the moving minimum ***e***(*K*(*t*), *ϵ*), the magenta trajectory reveals a large excursion. If *r*_*K*_ is low (F), the ball remains close to the moving stable state *e*(*K*(*t*), *ϵ*). As a result, the prey density *x* only shows slightly lower values than *e*_*x*_(*K*(*t*), *ϵ*) but they always remain above the extinction threshold *x*(*t*) > *x*_*e*_ (fig. 1E). By contrast as shown in fig. 1F, the large excursion of the ball causes the prey density *x* to drop temporarily below the extinction threshold *x*_*e*_ (gray dotted line). Hence, the transient dynamics dramatically change from persistence to a collapse of the prey population. This transition is solely triggered by the increase of the rate *r*_*K*_. Hence, the predator-prey system (1)–(2) exhibits R-tipping.

Since the rate of environmental change has to exceed a critical rate to cause R-tipping, the relation between different time scales – in particular between the time scale of environmental change and the time scale of the intrinsic ecosystem dynamics (the ‘reaction time’) – is a crucial determinant for this transition to occur (Wieczorek et al., 2011).

The relation between different time scales is also key to another ecological problem. Recent studies demonstrate that populations can withstand fast environmental change if they are able to respond comparatively fast by rapid evolution. Since, in these cases, populations which are maladapted to the newly established environmental conditions rescue themselves from going extinct, this phenomenon is called *evolutionary rescue* (Bell, 2013; Bell and Gonzalez, 2009; Carlson et al., 2014; Gomulkiewicz and Holt, 1995; Gomulkiewicz and Shaw, 2013; Gonzalez et al., 2013).

Most theoretical and empirical studies referring to evolutionary rescue focus on isolated populations that are exposed to environmental change (Gonzalez et al., 2013; Lachapelle and Bell, 2012; Martin et al., 2013; Ramsayer et al., 2013; Vander Wal et al., 2013). But naturally, whether populations are able to adapt to environmental change depends on their interactions with other species (Bastille-Rousseau et al., 2018; Lavergne et al., 2010; Lawrence et al., 2012; Schoener et al., 2001; Van der Putten et al., 2010). In fact, some studies have tackled evolutionary rescue in a multi-species context (De Mazancourt et al., 2008; Henriques and Osmond, 2020; Kovach-Orr and Fussmann, 2013; Norberg et al., 2012; Northfield and Ives, 2013; Osmond and de Mazancourt, 2013; Petkovic and Colegrave, 2019), some of which focused on predator-prey interactions (Cortez and Yamamichi, 2019; Osmond et al., 2017; Yamamichi et al., 2019; Yamamichi and Miner, 2015). In one of these works, Yamamichi and Miner (2015) demonstrate that rapid evolution of the prey alone can rescue its non-evolving predator from extinction. They call this mechanism *indirect evolutionary rescue*. The reverse situation - predators that are able to support prey persistence during environmental changes - is examined by Osmond et al. (2017). However, neither of the two does explicitly model environmental change by means of time-dependent parameters. Hence, both miss an essential point, namely that evolutionary rescue is the outcome of a race between rapid evolution and environmental change as suggested by (Alexander et al., 2014; Bell and Gonzalez, 2009; Carlson et al., 2014; Ferriere and Legendre, 2013; Gonzalez et al., 2013; Svensson and Connallon, 2019; Uecker et al., 2014).

In our work, we model this race against extinction by taking explicitly into account the dynamics of environmental change and the dynamics of evolution. To this end, we expose a prey population to a sufficiently fast environmental change at which it experiences a rate-induced collapse. Then we ask if a predator population is able to prevent this rate-induced collapse when it can adapt sufficiently fast to the declining prey population. Hence, our ultimate aim is to bring together rate-induced-tipping and indirect evolutionary rescue. Both processes are determined by a critical rate, when considered separately. However, combining both processes will lead to an expression which relates the critical rate of environmental change to the speed of evolution.

Furthermore, just like rate-induced tipping, (indirect) evolutionary rescue occurs on time scales which are comparable to the intrinsic dynamics of ecosystems. Hence, it can only be observed in the transients of eco-evolutionary systems. However, most theoretical studies analyzing eco-evolutionary systems solely focus on changes of the long-term behavior (Cortez and Ellner, 2010; Cortez and Patel, 2017; Yamamichi et al., 2011; Yoshida et al., 2003). Consequently, these studies neglect low population densities during the transient dynamics due to rate-induced tipping and therefore also miss out on recognizing counteracting processes such as evolutionary rescue (Hastings, 2001, 2004; Hastings et al., 2018; Vanselow et al., 2019). Our study overcomes this problem by focusing on the part of a time series which is essential for studying rate-induced tipping and evolutionary rescue, the transient dynamics.

The scope of our work can be outlined as follows: First, we sketch the idea of how rapid evolution of a predator’s trait can prevent a rate-induced prey collapse. We then introduce the complete eco-evolutionary predator-prey model. Based on this model, we demonstrate the occurrence of R-tipping in a scenario without evolution in which the trait is fixed to its minimum value. Afterwards, we show that rapid evolution of the predator’s trait can prevent this rate-induced collapse. In the following section, we demonstrate that the probability of rate-induced tipping decreases with increasing genetic variation. Finally, we consider the sensitivity of our results depending on the initial state of the system. In the last section, we discuss our results.

### How indirect evolutionary rescue can prevent rate-induced tipping

In this section, we study under which premises indirect evolutionary rescue is able to prevent the rate-induced collapse of the prey. Therefore, we add rapid evolution of the predator’s trait *α* as a third equation to the slow-fast predator-prey system (1)–(2). We model the trait dynamics (6) in accordance with the quantitative genetics approach derived in Lande (1982) and Abrams et al. (1993). In this framework, the rate of change of the mean trait value 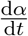 is proportional to the additive genetic variance of the predator population *r*_*V*_ *A*(*α*) and the fitness gradient 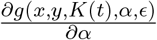 The three-dimensional eco-evolutionary system is given by:

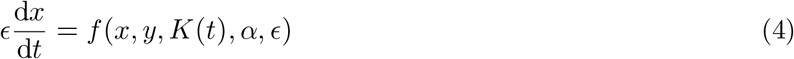

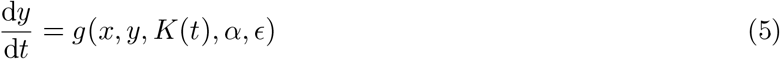

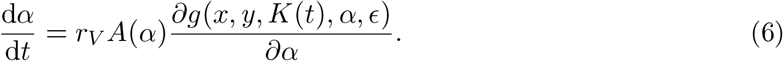

Again, we expose the eco-evolutionary system (4)–(6) to the same gradual environmental change *K*(*t*) = *K*_0_ − *r*_*K*_*t*, as given in Eq. (3), with *K*(*t*) ∈ [*K*_0_ *K*_*min*_] and assume that an unique moving stable state ***e***(*K*(*t*), *ϵ*) = (*e*_*x*_(*K*(*t*), *ϵ*), *e*_*y*_(*K*(*t*), *ϵ*), *e*_*α*_(*K*(*t*), *ϵ*)) exists with ***e***_*x*_(*K*(*t*), *ϵ*) > *x*_*e*_ for all values of *K*(*t*) ≥ *K*_*min*_. If the system (1)–(2) is able to stay close to the moving stable state during the transient, we say that it is tracking. For the eco-evolutionary system, this tracking means that the predator adjusts its trait *α* to the moving equilibrium value *e*_*α*_(*K*(*t*), *ϵ*) sufficiently fast to enable the coexistence of predator and prey *x, y* ≈ *e*_*x,y*_(*K, ϵ*). In the light of the gradual environmental degradation, we consider the case of perfect tracking (*x, y* = *e*_*x,y*_(*K, ϵ*)) as the optimal course of the eco-evolutionary system. Therefore, from now on, we refer to the moving stable equilibrium state ***e***(*K*(*t*), *ϵ*) as the optimum state ***e***_opt_(*K*(*t*), *ϵ*).

Whether the eco-evolutionary system can track the moving optimum state during the environmental change depends crucially on how fast *α* evolves in relation to the speed of environmental change *r*_*K*_. The trait *α* changes fast when the additive genetic variance *r*_*V*_ *A*(*α*) is high (Cortez, 2016, 2018). Notice that the additive genetic variance is the product of phenotypic variation *A*(*α*) and the genotypic or genetic variation expressed by the rate *r*_*V*_. However, in the following, we assume that the speed of evolution is mainly determined by genetic variation *r*_*V*_ (Carlson et al., 2014).

In figure 2, we analyze how the onset of the gradual environmental change of *K*(*t*) affects the dynamics of the eco-evolutionary system (4)–(6) which is initially situated in the optimum state ***e***_opt_(*K*(*t*), *ϵ*): *x*_0_ = *e*_opt,*x*_(*K*_0_, *ϵ*), *y*_0_ = *e*_opt,*y*_(*K*_0_, *ϵ*) and *α*_0_ = *e*_opt,*α*_(*K*_0_, *ϵ*). In the case of slow evolution (*r*_*V*_ is low in fig. 2A and fig. 2B), we again observe tracking if the rate *r*_*K*_ is low and R-tipping if the rate *r*_*K*_ is high. If the system is able to track the optimum state, the density of the prey population *x* always remains above the extinction threshold *x* > *x*_*e*_ (cyan curve) while it temporarily drops below the extinction threshold *x* < *x*_*e*_ between *t*_*e*_ and *t*_*r*_ when R-tipping occurs (magenta dashed line).

**Figure 2:**
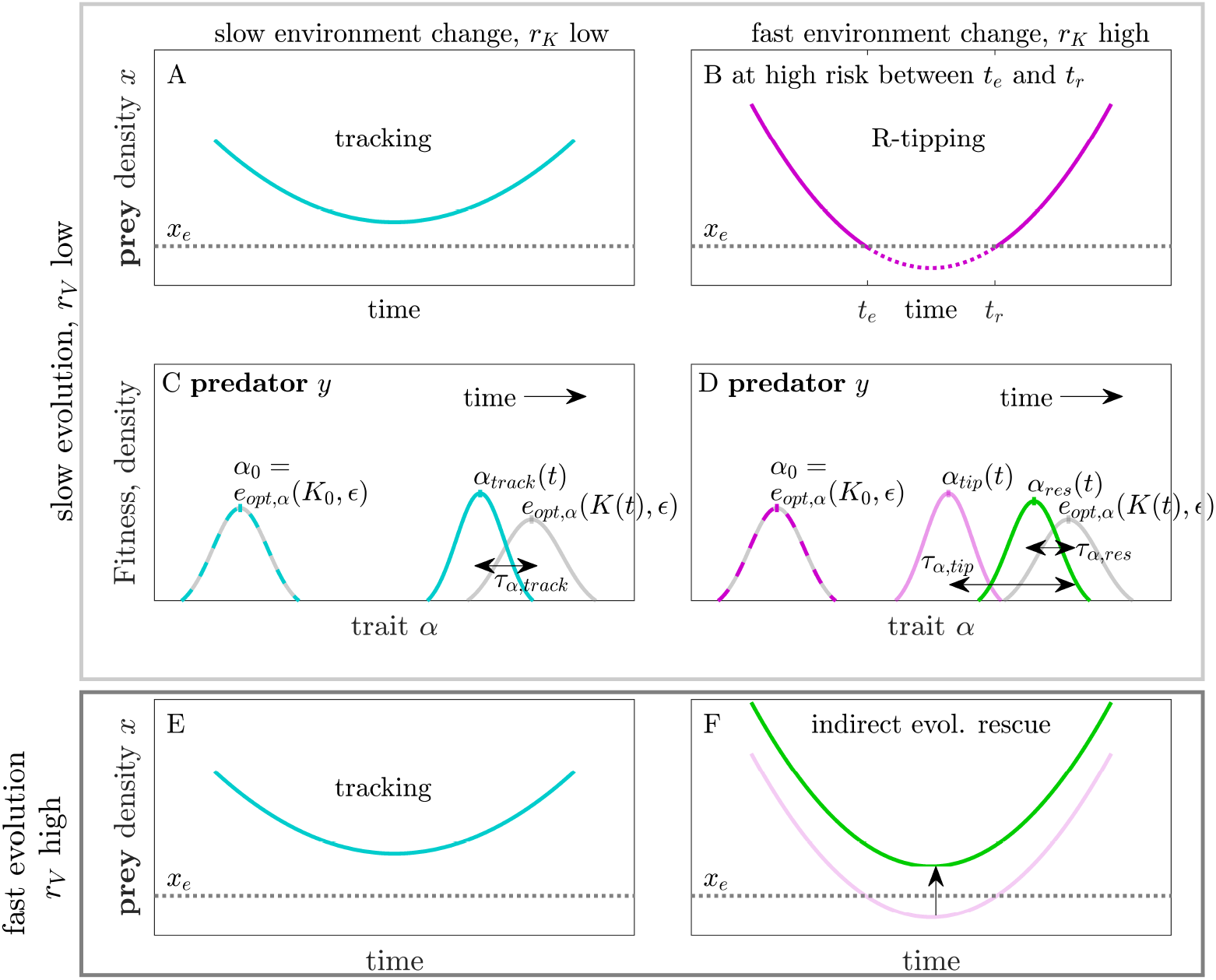
(A) The eco-evolutionary system (4)–(6) tracks: the prey population *x* declines but remains above the extinction threshold *x* > *x*_*e*_ when environmental conditions *K*(*t*) change slowly (*r*_*K*_ is low). (B) R-tipping: prey density *x* drops below the extinction threshold *x* < *x*_*e*_ between *t*_*e*_ and *t*_*r*_ because environmental conditions *K*(*t*) change too fast (*r*_*K*_ high). (C,D) Slow/fast decline of the prey causes small/large lag-load *τ*_*α*,track_/*τ*_*α*,tip_. (E) Distance between minimum prey density and extinction threshold *x*_*e*_ reduces when predator evolution is fast (*r*_*V*_ is high). (D,F) Indirect evolutionary rescue (green): the predator population can adapt faster (D) which lowers the lag-load: *τ*_*α*,res_ < *τ*_*α*,tip_ and prevents the rate-induced collapse of the prey (F).

As shown by figure 2C and D, this collapse can be explained by the emerging deviation of the mean trait value of the predator population *α*(*t*) from the optimum mean trait value *e*_opt,*α*_(*K*(*t*), *ϵ*). This deviation, which is called *lag-load τ*_*α*_(*K*(*t*), *ϵ*) = |*α*(*t*) − *e*_opt,*α*_(*K*(*t*), *ϵ*)| (Kopp and Matuszewski, 2014; Maynard Smith, 1976), quantifies the goodness of adaptation of the predator population: The smaller the lag-load *τ*_*α*_(*K*(*t*), *ϵ*), the better suited is the predator population to the current environmental conditions *K*(*t*).

Figure 2C and D demonstrate that the lag-load *τ*_*α*_(*K*(*t*), *ϵ*) is smaller in the case of tracking: *τ*_*α*_(*K*(*t*), *ϵ*) = *τ*_*α*,track_, and much larger in the case of R-tipping: *τ*_*α*,tip_ > *τ*_*α*,track_. Such a maladaption of the predator population provides a possible explanation of the rate-induced collapse of the prey. For instance, when the maladaption manifests in a predator population that is much more aggressive as it would be in the optimum *α*(*t*) > *e*_opt,*α*_(*K*(*t*), *ϵ*), then the prey is confronted with an unprecedented high predation pressure. This can result in prey population densities below the extinction threshold *x*_*e*_ (fig. 2B). By contrast, if the predator can adapt faster to the shrinking prey population (*r*_*V*_ is high), the lag-load *τ*_*α*,tip_ is reduced to *τ*_*α*,res_ < *τ*_*α*,tip_ (green curve in fig. 2D). In our example, this could lower the predation pressure on the prey and thus might prevent its rate-induced collapse (fig. 2F, green curve). In this case, indirect evolutionary rescue occurs.

### The eco-evolutionary predator-prey model

In the following, we demonstrate the interplay between rate-induced tipping and rapid evolution in a multi-species context using a paradigmatic ecosystem model. For this purpose, we replace the general equations (4)–(6) by the classical predator-prey system developed by Rosenzweig and MacArthur (Rosenzweig and MacArthur, 1963). In accordance with (Cortez and Ellner, 2010; Yamamichi et al., 2019), we neglect units and treat the eco-evolutionary system as a purely mathematical model:

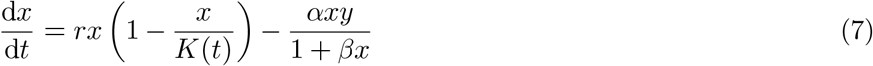

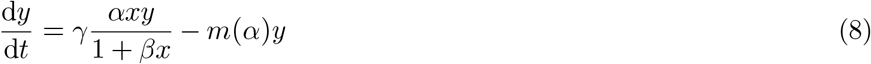

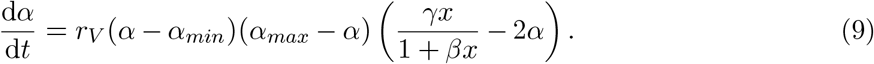

Here, the prey *x* grows logistically with maximum per–capita growth rate *r* and time-dependent carrying capacity *K*(*t*). Predator growth follows a Holling-Typ II functional response with maximum predation rate *α*/*β*, half-saturation constant 1/*β* and conversion efficiency *γ*. The predator’s mortality is assumed to be proportional to the mortality rate *m*(*α*) and the density of the predator *y*. The time scale separation between prey and predator *ϵ* is determined by the ratio between the predator’s mortality rate *m*(*α*) and the prey’s reproduction rate 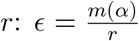. Because *ϵ* describes a ratio and is assumed to be 0 < *ϵ* < 1, the prey population always reproduces *ϵ*^−1^ times faster than its predator whose time scale is then given by 1.0.

In contrast to most other eco-evolutionary models, we explicitly model the temporal change of the environment by means of a continuously declining carrying capacity *K*(*t*) (Eq. (3)). The carrying capacity *K* is a suitable quantity as it defines the maximum prey density that the environment can ‘carry’. Accordingly, *K* is strongly linked to environmental conditions such as the availability of food sources, breeding sites or the suitability of climatic conditions. As discussed in the previous section, *K* is diminished at a rate *r*_*K*_ until a prescribed minimum *K*_*min*_ is reached. This minimum ensures that the prey does not go extinct due to the optimum falling below the extinction threshold. The realized scenario is comparable to a time-dependent degradation of the habitat due to climate variations, land use change or landscape fragmentation where the prey population loses access to essential resources.

In the eco-evolutionary model, the dynamical (evolutionary) trait of the predator is represented by its attack rate *α*. We assume that the attack rate *α* affects both predation 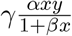 and mortality *m*(*α*) = (*α*^2^ + *c*)*y* of the predator. Therefore, an adaptation of *α* is associated with a trade-off: If the predator invests in its attack rate *α*, the predatory success will increase at the expense of a higher death rate (Abrams, 2000; Cortez and Ellner, 2010). In fact, it is this trade-off which enables a constructive adaptation of the predators attack rate depending on the available prey. Note that the trade-off crucially depends on how the attack rate *α* is incorporated into the mortality rate of the predator (see app. S3 for more details).

The dynamical equation for the attack rate *α* is derived according to Eq. (6). It is important to note that *α* depicts the mean trait within the predator population. Accordingly, the formulation of Eq. (9) is based on the assumption that the evolution of the mean trait is not significantly affected by the specific form of the frequency distribution of trait values within the predator population. By setting the phenotypic variance to *A*(*α*) = (*α* − *α*_*min*_)(*α*_*max*_ − *α*), we restrict the adaptation of *α* to an interval bounded by the lowest *α*_*min*_ and highest *α*_*max*_ reasonable value. The genetic variation *r*_*V*_ determines how fast the predator population is able to respond to changes (Cortez, 2018; Cortez and Weitz, 2014). Since evolutionary dynamics faster than the ecological dynamics are exceptional in natural systems (Cortez, 2018), we only consider evolutionary dynamics which are similar to or slower than the slowest time scale in the ecosystem - here predator reproduction. Therefore, we restrict our analysis to 0 < *r*_*V*_ ≤ 1.0.

Overall, the four-dimensional eco-evolutionary system (7)–(9) contains four different time scales: the time scale of the prey’s life cycle, the time scale of the predator’s life cycle expressed by their relationship 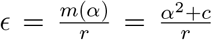, the rate of environmental change *r*_*K*_ and the speed of evolution represented by the genetic variation *r*_*V*_. The relationship between those time scales determines the dynamics of the system and will be the main focus of our study.

In the following, we discuss the dynamics of the eco-evolutionary system (7)–(9) using the exemplary parameter set given in table 1. Notice, that the rate of environmental change *r*_*K*_ and the speed of evolution *r*_*V*_ will be varied throughout our study.

**Table 1:** Parameters and initial conditions of the eco-evolutionary system (7)–(9). Furthermore, important bifurcations and terms.

| parameter | value |
| --- | --- |
| rate of environmental change $r_K$ | ***varies*** |
| speed of evolution $r_V$ | ***varies*** |
| growth rate $r$ | 0.9 |
| conversion efficiency $\gamma$ | 0.7 |
| handling efficiency $\beta$ | 0.4 |
| mortality $c$ | 0.1 |
| minimum attack rate $\alpha_{min}$ | 0.1 |
| maximum attack rate $\alpha_{max}$ | 1.3 |
| minimum carrying capacity $K_{min}$ | 1.0 |
| extinction threshold $x_e$ | $1/3 \cdot n_x(\alpha_0)$ |
| initial conditions | value |
| $x(0) = x_0$ | $n_x(\alpha_0)$ |
| $y(0) = y_0$ | $n_y(K_0, \alpha_0)$ |
| $K(0) = K_0$ | $K_T < K_0 < K_H$ |
| bifurcations and terms | value |
| $K_T(\alpha)$ (transcritical) | $\frac{\alpha^2+c}{\gamma\alpha-\alpha^2\beta-c\beta}$ |
| $K_H(\alpha)$ (Hopf) | $2K_T + \frac{1}{\beta}$ |
| $n_x(\alpha)$ | $\frac{\alpha^2+c}{\gamma\alpha-\alpha^2\beta-c\beta}$ |
| $n_y(K(t), \alpha)$ | $\frac{r}{\alpha} \left(1 - \frac{n_x}{K(t)}\right) (1 + \beta n_x)$ |

### Dynamics at the extreme trait values: no evolution but R-tipping

As has been discussed earlier, rate-induced tipping occurs as a system subjected to environmental change fails to track its moving stable state. In the context of eco-evolutionary dynamics, this moving stable state ***e***(*K*(*t*), *ϵ*) represents the evolutionary optimum state ***e***_opt_(*K*(*t*), *ϵ*) = (*e*_*x*,opt_(*K*(*t*), *ϵ*), *e*_*y*,opt_(*K*(*t*), *ϵ*), *e*_*α*,opt_(*K*(*t*), *ϵ*))) for the given environmental condition *K*(*t*). Accordingly, in order to examine how R-tipping occurs in the eco-evolutionary system (7)–(9), we first need to determine its time-dependent optimum state ***e***_opt_(*K*(*t*), *ϵ*).

We start by determining the optimum attack rate *e*_opt,*α*_(*K*(*t*), *ϵ*). Setting 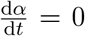 (9) reveals three different optimum attack rates *e*_opt,*α*_(*K*(*t*), *ϵ*): two given by the extreme trait values (i) *α* = *α*_*min*_ and (ii) *α* = *α*_*max*_ and one by the attack rate 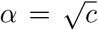 for which the fitness gradient vanishes (iii) 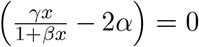. In this section, we concentrate on the two extreme cases which can only be maintained within the eco-evolutionary system if *α*(0) = *α*_0_ = *α*_*min*_ or *α*(0) = *α*_0_ = *α*_*max*_, respectively. Consequently, the trait *α*(*t*) remains constant for all time which ultimately excludes evolutionary adaptation. Though we are mainly interested in the case where evolution is active, it is quite instructive to demonstrate R-tipping without evolution first as it provides a benchmark for the following analyses.

As already stated, we only consider the dynamics for 0 < *ϵ* < 1.0. However, for *α* = *α*_*max*_, the time scale separation between predator and prey is given by 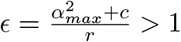 and thus the predator would reproduce faster than its prey. Consequently, we neglect this case and focus only on the case *α* = *e*_*α*,opt_(*K*(*t*), *ϵ*) = *α*_*min*_. In this scenario, the moving optimum ***e***_opt_(*K*(*t*), *ϵ*) can be written as:

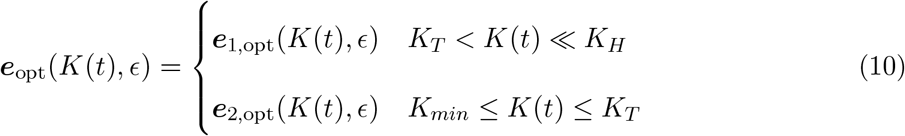

with ***e***_1,opt_(*K*(*t*), *ϵ*) = (*n*_*x*_(*α*_*min*_), *n*_*y*_(*K*(*t*), *α*_*min*_), *K*(*t*), *α*_*min*_) and ***e***_2,opt_(*K*(*t*), *ϵ*) = (*K*(*t*), 0, *K*(*t*), *α*_*min*_), where *n*_*x*_ and *n*_*y*_ fulfil 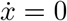 and 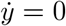 respectively (for *n*_*x*_, *n*_*y*_, *K*_*T*_ and *K*_*H*_ see tab. 1). For large values of *K*(*t*), a Hopf bifurcation occurs at *K*(*t*) = *K*_*H*_. However, in order to avoid long-term oscillations, we choose *K*_0_ ≪ *K*_*H*_. The critical value *K*_*T*_ marks the transcritical bifurcation at which ***e***_1,opt_(*K*(*t*), *ϵ*) loses stability, while ***e***_2,opt_(*K*(*t*), *ϵ*) becomes stable.

In figure 3, we illustrate how the optimum prey ***e***_opt,*x*_(*K*(*t*), *ϵ*) (A) and predator densities ***e***_opt,*y*_(*K*(*t*), *ϵ*) (B) change in time (gray dashed lines) due to their dependence on the time-varying parameter *K*(*t*). We find a constant optimum prey and a decreasing optimum predator population density as long as *K*(*t*) > *K*_*T*_ (vertical dotted magenta line). Beyond the transcritical bifurcation *K*_*T*_ the predator is extinct in the moving optimum state, while the optimum prey density decreases linearly in time as ***e***_opt,*x*_(*K*(*t*), *ϵ*) = *K*(*t*) until *K*_*min*_ is reached (horizontal magenta dotted line). Afterwards, the optimum prey density remains constant at the long-term state ***e***_opt,*x*_(*K*_*min*_, *ϵ*) = *K*_*min*_. As *K*(*t*) decreases linearly in time, we expect the prey population to persist and the predator to go extinct if the eco-evolutionary system (7)–(9) is able to track the moving optimum state ***e***_opt_(*K*(*t*), *ϵ*) during the transient.

**Figure 3:**
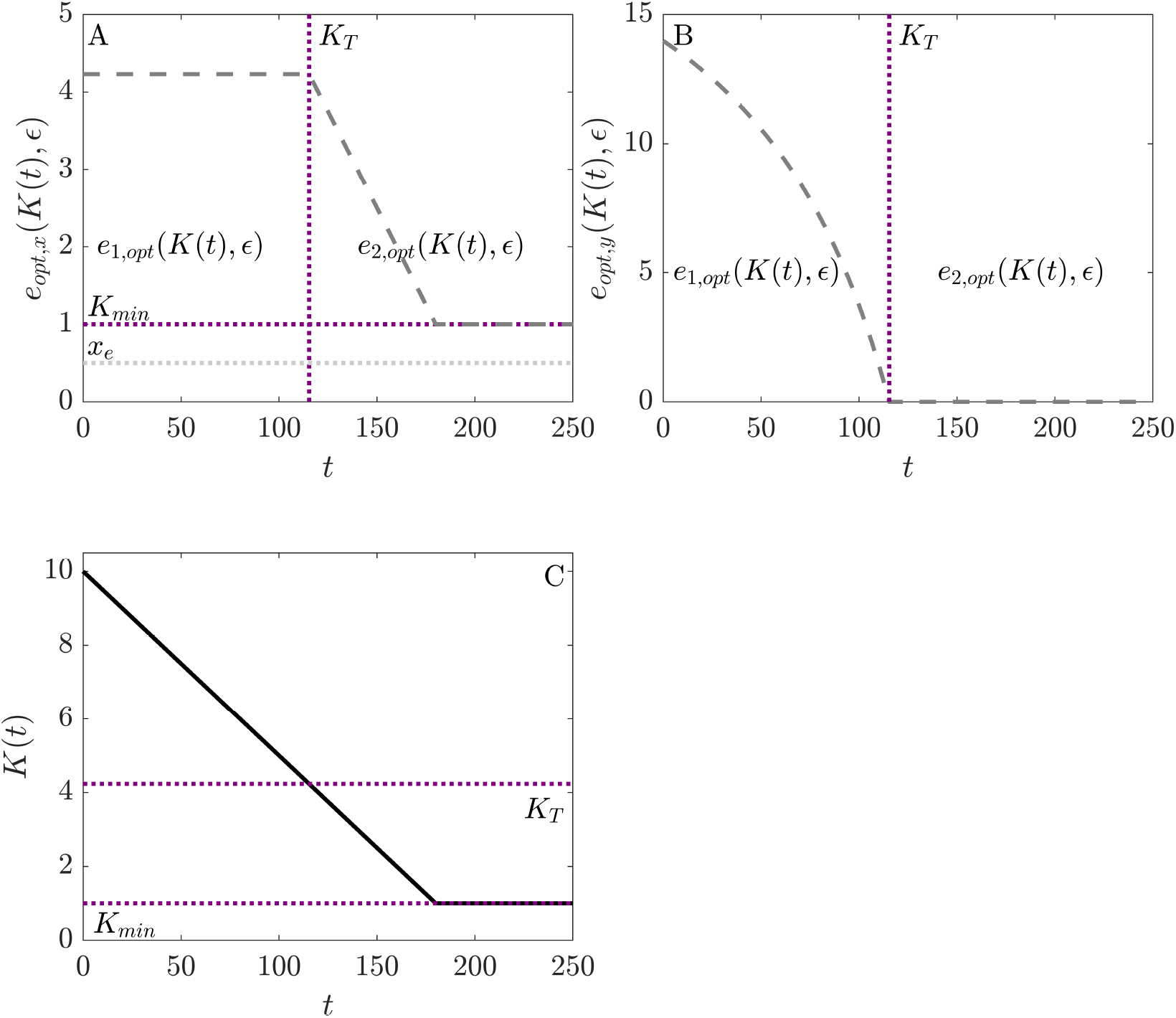
How optimum prey ***e***_opt,*x*_(*K*(*t*), *ϵ*) (A) and predator density ***e***_opt,*y*_(*K*(*t*), *ϵ*) (B) (gray dashed line) change in time. For *K*(*t*) > *K*_*T*_, *e*_1,opt_(*K*(*t*), *ϵ*) represents the moving optimum state. At *K*_*T*_ (vertical magenta dotted line), *e*_1,opt_(*K*(*t*), *ϵ*) and *e*_2,opt_(*K*(*t*), *ϵ*) exchange stability in a transcritical bifurcation. For *K*(*t*) < *K*_*T*_, *e*_2,opt_(*K*(*t*), *ϵ*) is the moving optimum state. (C) Linear decline of the carrying capacity *K*(*t*) at the rate *r*_*K*_ according to (3). Parameter: *x*_0_ = *n*_*x*_(*α*_*min*_), *y*_0_ = *n*_*y*_(*K*(*t*), *α*_*min*_), *K*_0_ = 10.0, *α*_0_ = *α*_*min*_, *r*_*K*_ = 0.05, *r*_*V*_ = 0. Other parameters see tab. 1.

As depicted in figure 4, we now study the transient dynamics of the eco-evolutionary system (7)– (9) for the same initial condition ***e***_opt_(*K*_0_ = 10, *α*_0_ = *α*_*min*_) (10) but different rates of environmental change *r*_*K*_. For a small rate of environmental change *r*_*K*_ = 0.05 (A), we obtain the expected behavior, namely that the system is able to track the moving optimum (gray dashed line). As a consequence, the prey density *x* always remains above the extinction threshold *x*_*e*_ while the predator population goes extinct in the long run. Interestingly, its extinction occurs much later than expected from the optimum predator density ***e***_opt,*y*_(*K*(*t*), *ϵ*) (gray dashed line).

**Figure 4:**
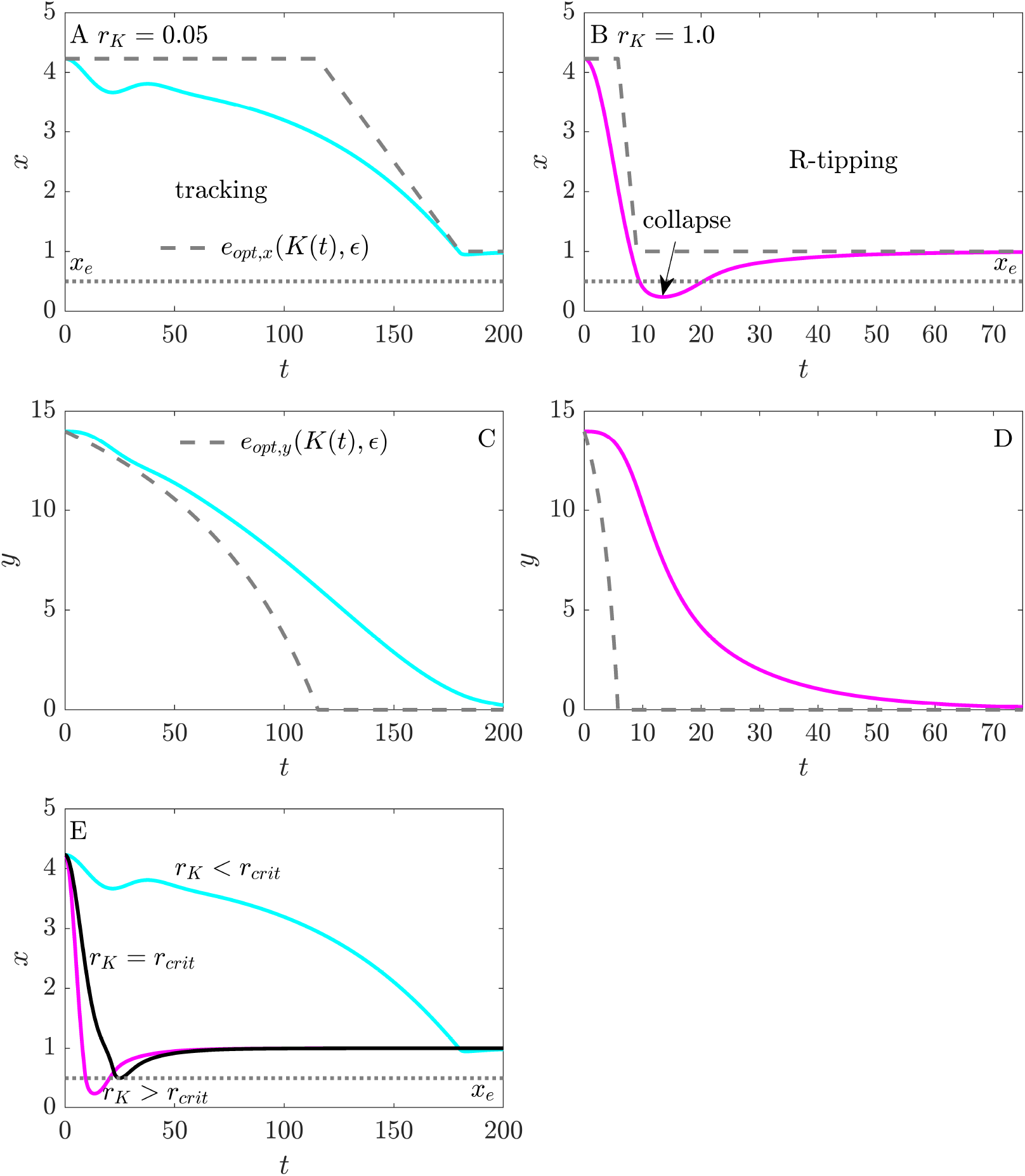
(A,B) The eco-evolutionary system (7)–(9) shows tracking (cyan) of the optimum state ***e***_opt_(*K*(*t*), *ϵ*) (gray dashed line) at low rate *r*_*K*_ = 0.05: prey density remains above extinction threshold *x*_*e*_ (gray dotted line). (C,D) System shows R-tipping (magenta) at higher rate *r*_*K*_ = 1.0: prey density drops below extinction threshold *x*_*e*_. (E) The critical rate *r* = *r*_*crit*_ (black) is defined as the lowest rate *r*_*K*_ at which *x* ≤ *x*_*e*_ during the transient dynamics. The system tracks/tips for *r*_*K*_ < *r*_*crit*_/*r*_*K*_ > *r*_*crit*_. Parameter: *x*_0_ = *n*_*x*_(*α*_*min*_), *n*_*y*_(*K*_0_, *α*_*min*_), *K*_0_ = 10.0, *α*_0_ = *α*_*min*_ and tab. 1.

For a higher rate *r*_*K*_ = 1.0 (fig. 4B), we observe the rate-induced collapse of the prey. This collapse occurs because a small prey population, which constantly holds densities below its optimum *x* < ***e***_opt,*x*_(*K*(*t*), *ϵ*), is confronted with a predator whose density is above its optimum *y*> ***e***_opt,*y*_(*K*(*t*), *ϵ*). The resulting overwhelming predation pressure ultimately forces the prey population to drop below the extinction threshold *x*_*e*_.

To clearly separate tracking *r*_*K*_ < *r*_*crit*_ from R-tipping *r*_*K*_ > *r*_*crit*_, we define the critical rate *r*_*crit*_ as the lowest rate *r*_*K*_ at which *x* ≤ *x*_*e*_ during the transient dynamics. It should be noted that this definition is fundamentally different from the mathematical definition of R-tipping as outlined in (Ashwin et al., 2012; Vanselow et al., 2019; Wieczorek et al., 2011).

In the next section, we study whether we still find a rate-induced collapse of the prey if the predator population is capable of evolutionary adaptation. In such a case, the predator might be able to reduce the distance to the optimum state which would lower the predation pressure on the prey population and thereby possibly prevent the collapse.

### Indirect evolutionary rescue prevents rate-induced tipping

In the following, we study whether a sufficiently fast evolutionary adaptation of the predator population can prevent the rate-induced collapse of the prey. Therefore, we choose an initial attack rate *α*_0_ between the two extreme values *α*_*min*_ and *α*_*max*_ and study the dynamics of the eco-evolutionary system (7)–(9) depending on the rate of environmental change *r*_*K*_ and the genetic variation *r*_*V*_. In this case, the eco-evolutionary system (7)–(9) possesses the moving optimum state ***e***_opt_(*K*(*t*), *ϵ*):

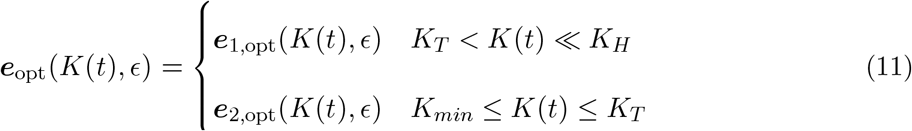

with 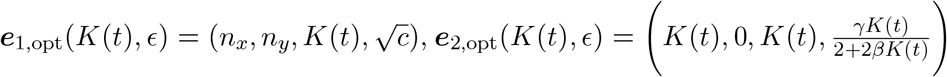 and 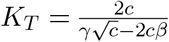 respectively 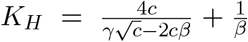 (see tab. 1 for *n*_*x*_, *n*_*y*_ and other parameters). At *K*(*t*) = *K*_*T*_, the system passes a transcritical bifurcation where ***e***_1,opt_(*K*(*t*), *ϵ*) and ***e***_2,opt_(*K*(*t*), *ϵ*) exchange stability (see fig. 3).

When *r*_*V*_ = 0.1, as expected, increasing the rate of environmental change from *r*_*K*_ = 0.05 (fig. 5A) to *r*_*K*_ = 1.0 (fig. 5B) causes the rate-induced collapse of the prey. In the latter case the prey population is confronted with an extraordinary predation pressure because the attack rate very quickly becomes much higher than the optimum attack rate: *α* > ***e***_opt,*α*_(*K*(*t*), *ϵ*) (fig. 5K). As a consequence, the *signed lag-load* 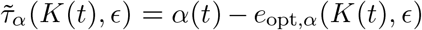 is large and positive (fig. 5K). Note, that we show the *signed lag-load* 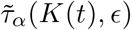 instead of the usually applied absolute *lag-load τ*_*α*_ (Gomulkiewicz and Holt, 1995) *- to emphasize the importance of the direction of maladaptation which is reflected by the sign of* 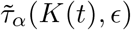. But as shown in figure 5C, this rate-induced collapse can be prevented if the genetic variation is high enough, here *r*_*V*_ = 0.9 instead of *r*_*V*_ = 0.1. The faster adaptation of the predator population suppresses the high initial attack rates. Instead, the attack rate *α* drops rather rapidly and attains rather low values - compared to the optimum and the scenarios representing tracking and R-tipping - before it approaches slowly the optimum. The mostly negative signed lag-load 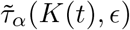 (fig. 5) means that predators are less aggressive than in the optimum for most of the time which lowers the predation pressure on the prey and prevents its rate-induced collapse. As it is the predator whose adaptation secures the prey’s survival, it is a case of indirect evolutionary rescue.

**Figure 5:**
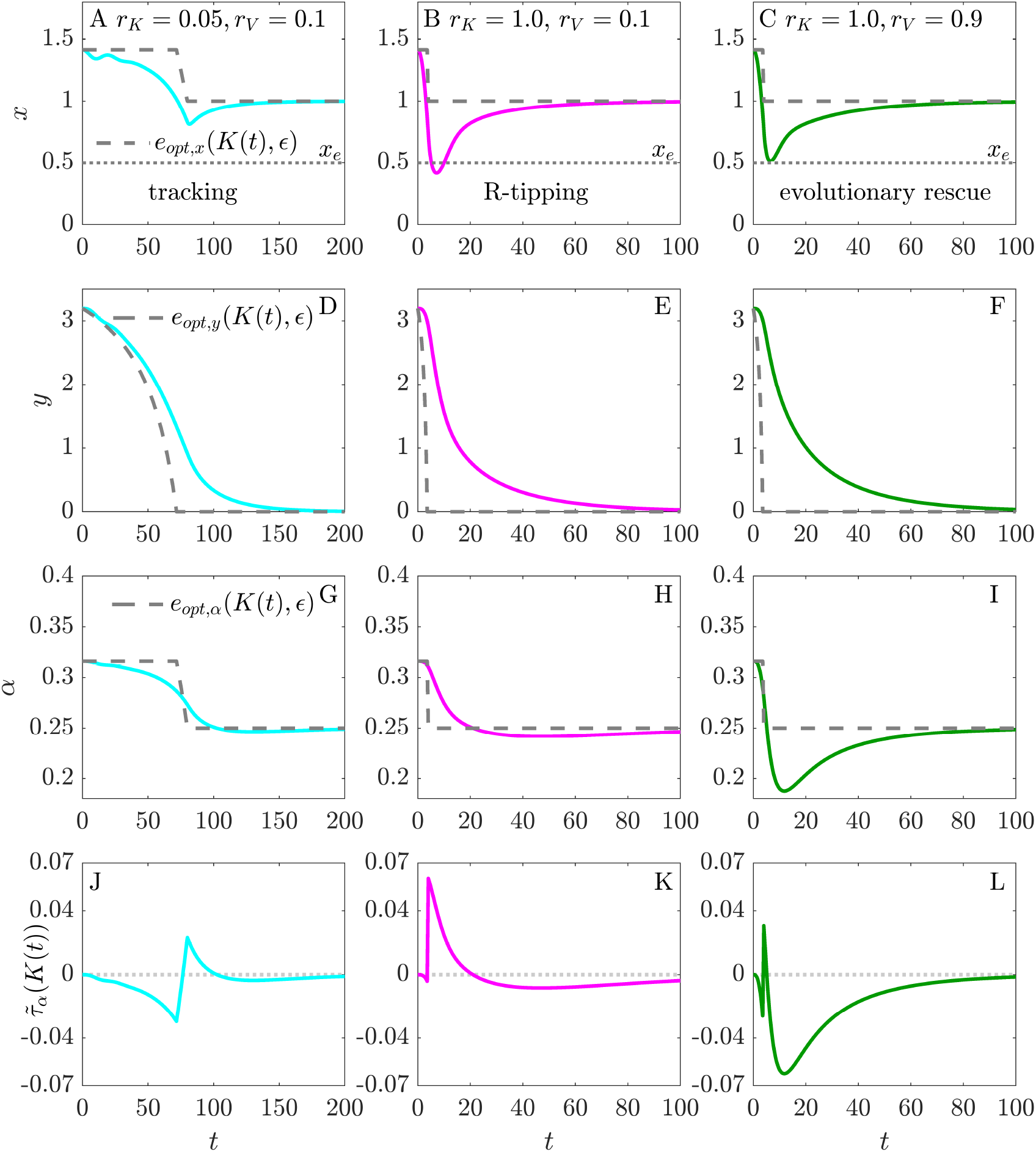
(A, B, C) Prey density *x*, (D, E, F) predator density *y*, (G, H, I) attack rate *α* of the eco-evolutionary system (7)–(9) and (J,K,L) signed lag-load 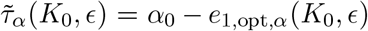 for different combination of the rate of environmental change *r*_*K*_ and the rate of evolution *r*_*V*_. The left column shows tracking (cyan), the middle column depicts R-tipping (magenta) and the right column demonstrate indirect evolutionary rescue (green). Parameters: *x*_0_ = *e*_opt,*x*_(*K*_0_, *ϵ*), *y*_0_ = *e*_opt,*y*_(*K*_0_, *ϵ*), *K*_0_ = 5.0, *α*_0_ = *e*_opt,*α*_(*K*_0_, *ϵ*), other parameters see tab. 1.

In summary, we find three different transient dynamics: tracking, R-tipping and indirect evolutionary rescue depending on the relation of the rate of environmental change *r*_*K*_ and the genetic variation *r*_*V*_. In the next section, we evaluate this relationship in more detail.

### The probability of rate-induced tipping decreases with increasing genetic variation

To obtain a comprehensive picture of rate-induced tipping and its prevention by indirect evolutionary rescue, we estimate the critical rate of environmental change *r*_*crit*_ depending on the rate of evolutionary adaptation *r*_*V*_. This critical rate separates the two possible outcomes, R-tipping (magenta region in fig. 6) or tracking (cyan and green regions). As a certain rate of environmental change has to be exceeded to induce R-tipping, without evolutionary adaptation, the tracking region can be divided into two parts: One where evolutionary rescue is not necessary to obtain tracking (cyan) and one where it is (green). The upper limit of the latter is depicted by the critical rate *r*_*crit*_ (black curve) above which R-tipping occurs. The critical rate portrays the expected relationship: the higher the genetic variation *r*_*V*_, the lower the probability of R-tipping. Accordingly, a predator population with higher genetic variation is able to reduce its attack rate sufficiently fast to balance the gradual degradation of the prey’s carrying capacity (decrease of *K*(*t*)) and thus prevent its rate-induced collapse.

**Figure 6:**
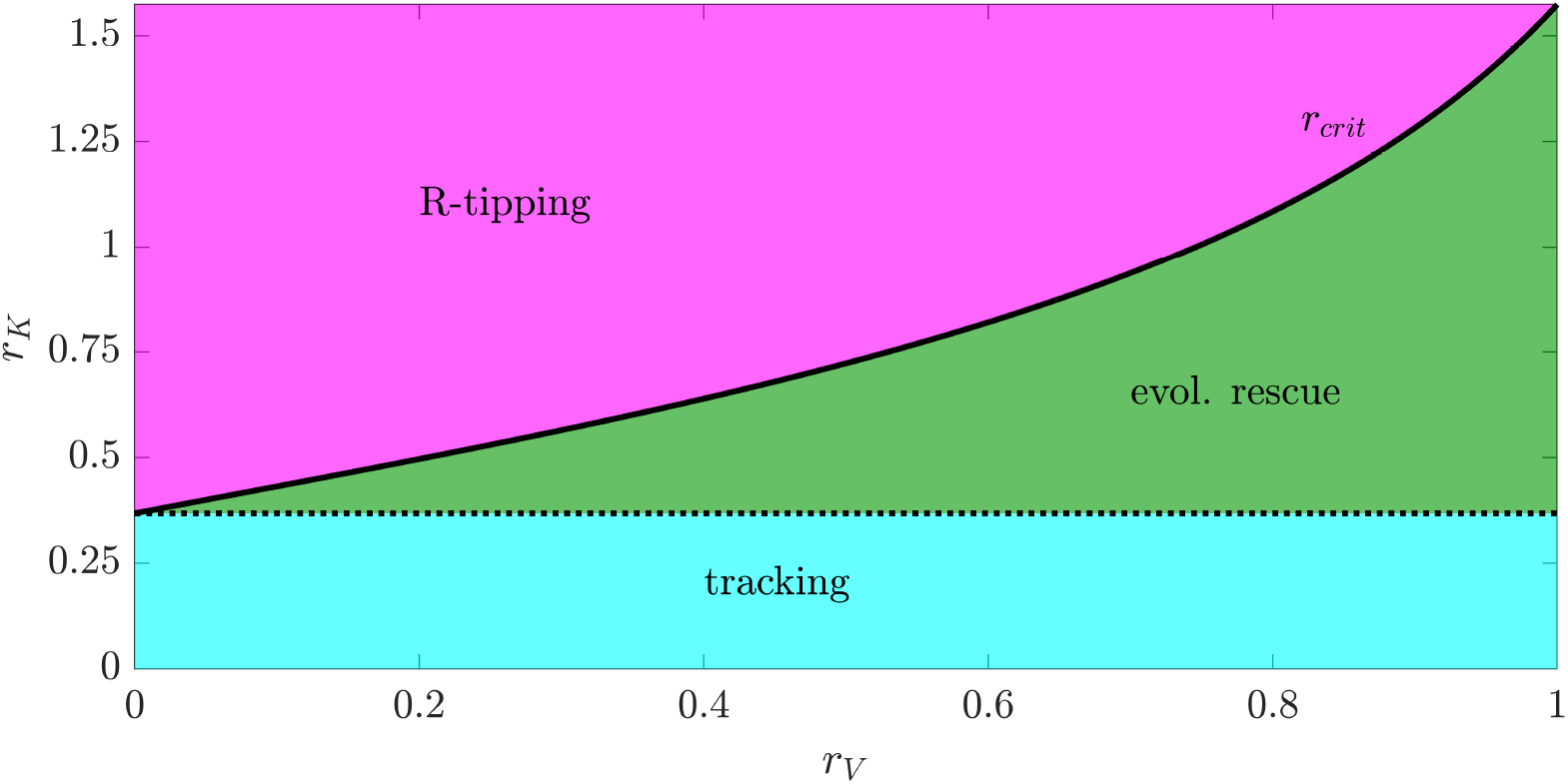
Critical rate *r*_*crit*_ (black curve) increases with rate of evolution *r*_*V*_. It separates the (*r*_*K*_, *r*_*V*_)-parameter space into the R-tipping region (magenta) and the tracking-region (green/cyan). The black dotted line further divides the tracking-region into the area where R-tipping is impossible (cyan) and the area where indirect evolutionary rescue enables tracking (green). Parameter *K*_0_ = 5.0, *α*_0_ = 0.6, *x*_0_ = *n*_*x*_(*α*_0_), *y*_0_ = *n*_*y*_(*α*_0_, *K*_0_). Other parameters see tab. 1.

### The occurrence of indirect evolutionary rescue varies with initial conditions

Hereinafter, we examine if the appearance of R-tipping or indirect evolutionary rescue depends on the initial conditions of the eco-evolutionary system (7)–(9). So far, we have assumed that the system is always situated in the optimum state in the beginning. As a result, the predator population is optimally adapted before the carrying capacity starts to change. Consequently, the initial lag-load is given by *τ*_*α*_(*K*_0_, *ϵ*) = 0 as illustrated in figure 2. In the following, we vary the initial attack rate *α*_0_ and the initial value of the carrying capacity *K*_0_ independently of one another. Hence, the predator population becomes less well adapted to the initial environmental condition *K*_0_. To further ensure a reasonable relation between the initial prey and predator density *x*_0_ and *y*_0_, we set *x*_0_ and *y*_0_ to the optimum equilibrium densities *x*_0_ = *e*_1,opt,*x*_(*K*_0_, *ϵ*) = *n*_*x*_(*α*_0_) and *y*_0_ = *e*_1,opt,*y*_(*K*_0_, *ϵ*) = *n*_*y*_(*K*_0_, *α*_0_) (see tab 1 for *n*_*x*_(*α*) and *n*_*y*_(*K*(*t*), *α*)). Hence, the initial conditions (*n*_*x*_(*α*_0_), *n*_*y*_(*K*_0_, *α*_0_), *K*_0_, *α*_0_) are fully determined by setting the initial attack rate *α*_0_ and the initial carrying capacity *K*_0_.

At first, we study exemplary initial conditions with different initial attack rates *α*_0_ ≷ *e*_opt,*α*_(*K*_0_, *ϵ*) (fig. 7). For *α*_0_ < *e*_opt,*α*_(*K*_0_, *ϵ*), the higher the initial carrying capacity *K*_0_ = 4.5 and the genetic variation *r*_*V*_ = 0.9, the closer is the minimum of the prey density to the extinction threshold *x*_*e*_ (fig. 7A). For *α*_0_ > *e*_opt,*α*_(*K*_0_, *ϵ*), the prey density shows the opposite behavior: instead of dropping, it increases in the beginning and reveals the highest distance to the extinction threshold for high values of *K*_0_ and *r*_*V*_, *K*_0_ = 4.5 and *r*_*V*_ = 0.9 (fig. 7B). This contradiction can be explained by studying the initial densities of the predator population (fig. 7C,D) and the change of the attack rate *α* (fig. 7E,F). For *α*_0_ < *e*_opt,*α*_(*K*_0_, *ϵ*), initial predator densities are consistently high while the attack rate increases. Consequently, the predation pressure increases leading to the early drop of the prey population density (fig. 7G). Conversely for *α*_0_ > *e*_opt,*α*_(*K*_0_, *ϵ*), initial densities of the predator population are much lower and the attack rate decreases. This reduces the predation pressure (fig. 7H) causing the temporary increase of the prey density in the beginning. Hence, we expect prey populations with higher initial carrying capacity *K*_0_ exposed to predators with *α*_0_ > *e*_opt,*α*_(*K*_0_, *ϵ*) and higher genetic variation *r*_*V*_ to be most likely to experience indirect evolutionary rescue.

**Figure 7:**
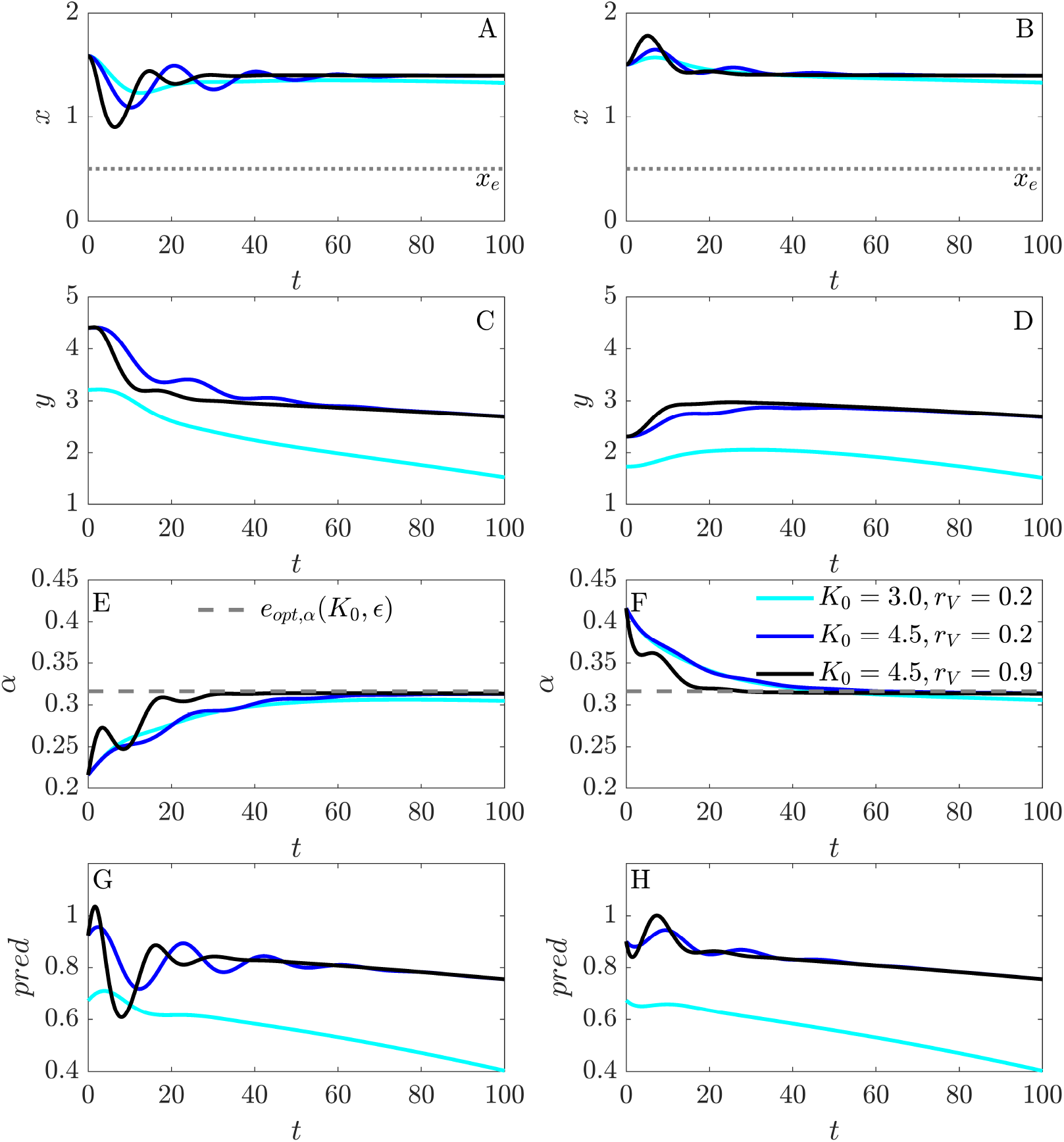
(A,B) Prey density *x*, (C,D) predator density *y*, (E,F) attack rate *α* and (G,H) predation pressure 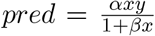 for *α*_0_ < *e*_opt,*α*_ (left column) and *α*_0_ > *e*_opt,*α*_ (right column). Parameters: *α*_0_ = *e*_1,opt,*α*_ ± 0.1, *x*_0_ = *n*_*x*_(*α*_0_), *y*_0_ = *n*_*y*_(*K*_0_, *α*_0_), *r*_*K*_ = 0.1. Other parameters tab. 1.

In figure 8, we now study the occurrence of evolutionary rescue for low (A, *r*_*V*_ = 0.2) and high genetic variation (B, *r*_*V*_ = 0.9) for initial conditions *K*_0_ ∈ [3, 5] and *α*_0_ ∈ [0.1, 0.5]. We choose these intervals to ensure the presence of the predator population and the absence of long-term oscillations. Therefore, initial conditions located in the gray regions are not taken into account.

**Figure 8:**
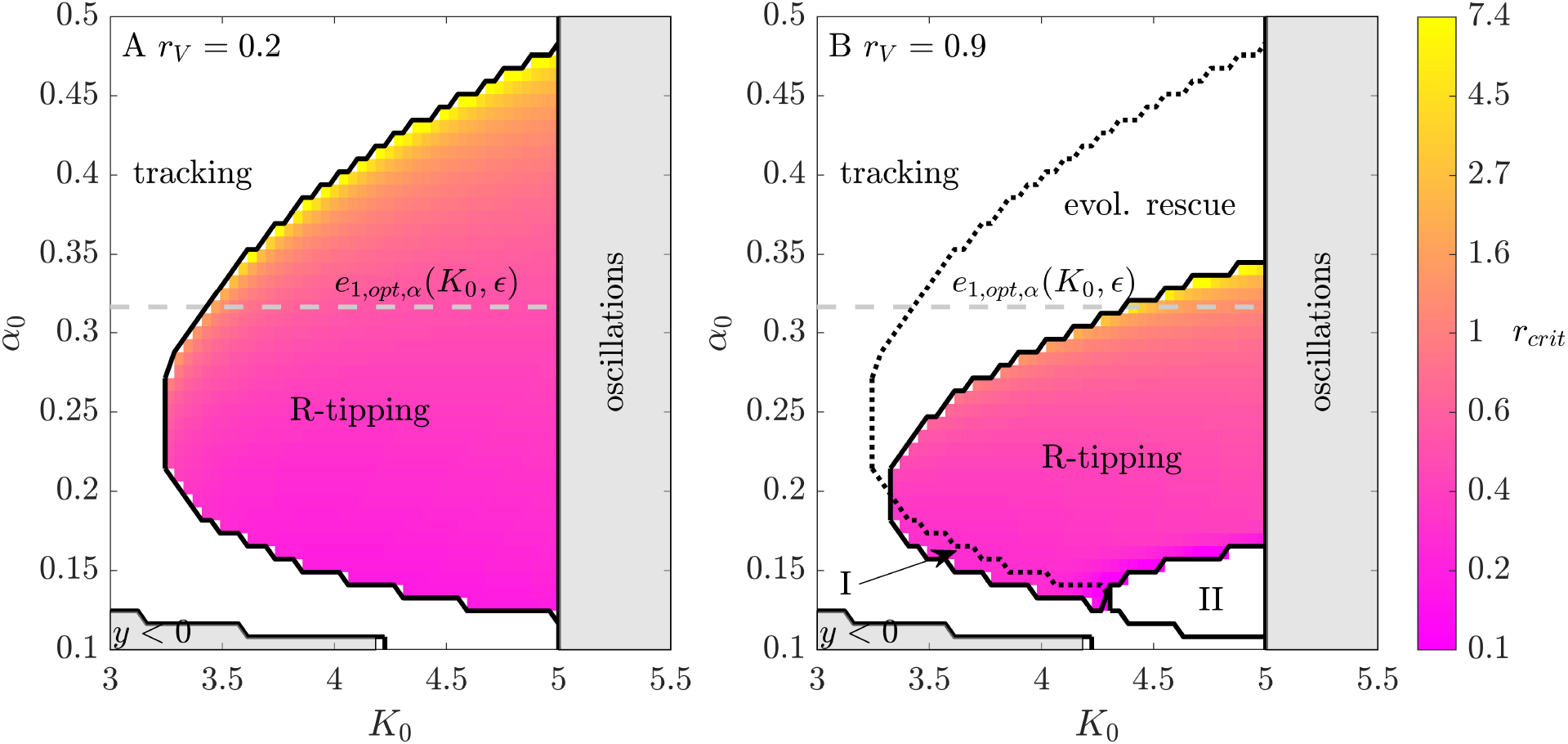
(*K*_0_, *α*_0_)-parameter space for *r*_*V*_ = 0.2 (A) and *r*_*V*_ = 0.9 (B). Initial conditions in the coloured-region show R-tipping above the critical rate *r*_*crit*_ (yellow: high, magenta: low). Initial conditions in the white non-coloured region track. Initial conditions showing long-term oscillations or negative initial predator densities are not considered (gray regions). Initial conditions determined by higher initial attack rates experience indirect evolutionary rescue while initial conditions with lower initial attack rates change its behavior from tracking to R-tipping (coloured-region I). Initial conditions in II already achieve prey densities *x* < *x*_*e*_ due to transient oscillations if *r*_*K*_ = 0. Other parameters see tab. 1.

In figure 8, the critical rate *r*_*crit*_ of environmental change increases as the initial attack rate *α*_0_ achieves higher values. Accordingly, initial conditions with *α*_0_ > *e*_opt,*α*_(*K*_0_, *ϵ*) are less prone to exhibit R-tipping. In this case, the prey population experiences lower predation pressure because (i) initial predator densities are lower and (ii) the attack rate decreases sufficiently fast (see fig. 7F). As expected, these initial conditions experience indirect evolutionary rescue when the genetic variation is increased from *r*_*V*_ = 0.2 (A) to *r*_*V*_ = 0.9 (B). In the latter case, the already low predation pressure can be reduced even further due to a faster decreasing attack rate *α*.

Interestingly, some initial conditions which are located in the tracking region for *r*_*V*_ = 0.2 exhibit R-tipping as *r*_*V*_ is increased to 0.9 (region I in fig. 8B). This means that they tip precisely because of the evolutionary adaptation (increasing attack rate) being too fast which ultimately causes a higher predation pressure than for *r*_*V*_ = 0.2 (compare fig. 7G). A rather extreme case of this mechanism can be found in region II where the prey density has already dropped to densities below *x*_*e*_ during the transient oscillation when *r*_*K*_ = 0 (see fig. 7 for the transient oscillation).

Furthermore, figure 8 demonstrates that the same initial lag-load *τ*_*α*_(*K*_0_, *ϵ*) = |*α*_0_ − *e*_1,opt,*α*_(*K*_0_, *ϵ*)| can result in two completely different and contrasting outcomes: the occurrence of indirect evolutionary rescue or the rate-induced collapse of the prey population. This again emphasizes that aside from the magnitude of maladaptation, the direction of initial maladaptation - the signed lag-load 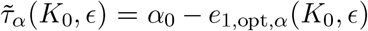 - can be a guiding quantity when studying the sensitivity of populations to the sudden onset of fast environmental changes.

## Discussion

### The relation of environmental and evolutionary time scale

We have demonstrated that rapid evolution of a predator’s trait can prevent a rate-induced collapse of its prey. Whether this indirect evolutionary rescue is put into effect depends crucially on the relation between the rate of environmental change and the rate of evolutionary adaptation. We highlighted this relationship by deriving a critical rate of environmental change, above which R-tipping can be expected, depending on the genetic variation which determines the rate of evolution (see fig. 6). We found that R-tipping is less and thus indirect evolutionary rescue is more likely when the genetic variation is high. This finding aligns well with former studies. For instance, Yamamichi and Miner (2015) who, however, did not explicitly consider rates of environmental change found that a higher genetic variation of a prey population shortens the vulnerable period of its non-evolving predator. Further theoretical (Barrett and Schluter, 2008; Gomulkiewicz and Holt, 1995; Vander Wal et al., 2013) and empirical studies (Ramsayer et al., 2013) show that high genetic variation increases the probability of evolutionary rescue of isolated populations.

As the rates of environmental change and evolutionary adaptation are of paramount importance for the outcome environmental changes, it is useful to consider under which circumstances they might be altered. For instance, populations might become more prone to R-tipping in the future as the speed of environmental change will increase due to climate change (Jezkova and Wiens, 2016; Nadeau and Urban, 2019; Shefferson et al., 2017). As suggested by the temporary change of the carrying capacity (fig. 3C), even short-term changes of environmental conditions can potentially provide an adequate explanation for an observed population collapse. On the contrary, recent studies suggest that this trend could be counteracted by the dispersal of sub-populations between different patches (Arumugam et al., 2020; Bourne et al., 2014). As shown by Bourne et al. (2014), the underlying mechanism is that dispersal can increase the genetic variation of populations when beneficial mutations are spread across sub-populations. Furthermore, the recent study by Arumugam et al. (2020) demonstrates that even when evolutionary adaptation is neglected, the dispersal between predator-prey populations reduces the probability of R-tipping. However, this mechanism requires the existence of patches as well as a rather large overall population size. Furthermore, a loss of genetic variation occurs naturally if a population suddenly shrinks (Baden et al., 2019; Fountain et al., 2016). Even though, if this decline was transient, such populations would be more susceptible to rate-induced tipping in the following as its genetic variability would have already been degraded. Such maladaptations can ultimately lead to the extinction of a population but can also affect other populations which they interact with (i.e. their prey) (Osmond et al., 2017; Yamamichi et al., 2019; Yamamichi and Miner, 2015).

Finally it should be noted that the specific relation between the rate of environmental change and the genetic variation is strongly linked to the choice of the extinction threshold. We introduced the extinction threshold in order to portray a population’s high endangerment when it undergoes stages of low density during its transient. One possibility to transfer this into the long-term behavior of the model would be the inclusion of the Allee effect in the dynamics of the prey (Allee et al., 1949). The Allee effect is useful in the context of eco-evolutionary systems because it incorporates a correlation between fitness and population size (Courchamp et al., 1999; Stephens and Sutherland, 1999; Stephens et al., 1999).

### Rapid evolution can stabilize or destabilize the transient dynamics

Furthermore, we have found that a sufficiently fast increase of the predator’s attack rate can cause indirect evolutionary rescue whereas its fast decline can lead to R-tipping (fig. 7 and fig. 8). Hence, rapid evolution of the predator’s trait can stabilize (by preventing low densities) or destabilize (by causing low densities) the transient state of the prey. Stabilizing and destabilizing evolution have so far been associated with the replacement of long-term oscillations by long-term stationary coexistence of populations and vice versa (Abrams, 2000; Cortez and Patel, 2017; Cortez et al., 2020; Mougi and Iwasa, 2010). In this context, (indirect) evolutionary rescue and R-tipping can be seen as additional mechanisms which are able to stabilize or destabilize, respectively, the state of populations on ecologically relevant (short) time scales (during the transient). Interestingly, this stabilizing (destabilizing) effect of rapid evolution is independent of the long-term persistence of the predator population (compare fig. 5 and fig. S4). This again underlines the importance of considering transient states of populations instead of analyzing exclusively their long-term behavior, as has already been emphasized by Hastings et al. (2018).

## Conclusion

All of our findings become solely visible in the transient dynamics of an eco-evolutionary model. Clearly, commonly applied models are an oversimplification of reality but as shown here, they can give valuable insights into the interaction of counteracting processes on ecologically relevant time scales. Our results show that we have to explicitly include rates of environmental change when modelling the race between evolutionary adaptation and extinction. Moreover, we suggest that besides the degree of maladaption (lag-load), its direction (signed lag-load) – trait lower/higher than optimum – is a decisive factor in inducing R-tipping or indirect evolutionary rescue. Finally, we once again emphasize the necessity of considering the transient dynamics because solely the transient reveals important processes on short time scales such as R-tipping or evolutionary rescue. Unraveling the mechanisms behind these processes will help us to understand the response of populations to habitat destruction or fragmentation and recent as well as future climate change.

## S1 Time-scale separation

To study the time-scale separation between predator and prey, we rewrite the predator-prey system (S1)–(S2) in dimensionless prey density *u*, predator density *v* and dimensionless time 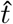:

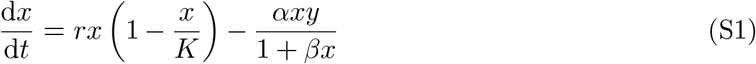

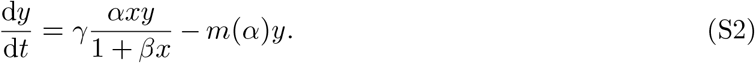

With 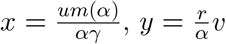 and 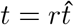, the predator-prey system can be written as:

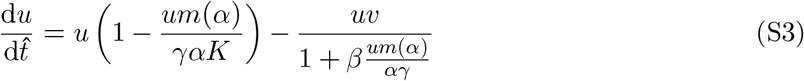

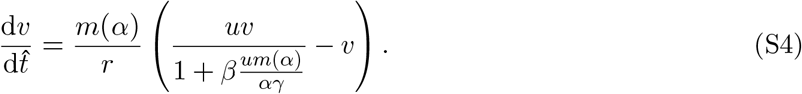

By setting 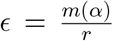 as time scale parameter, we can write the dimensionless predator-prey system in slow time 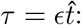:

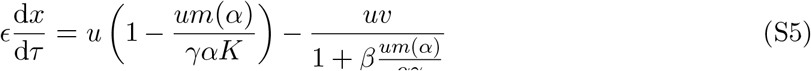

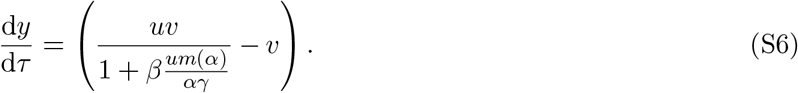

For *m*(*α*) = *α*^2^ + *c*, the separation between the prey’s and predator’s time scale is given by 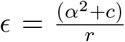. When *α*^2^ + *c* = *r*, prey and predator evolve on the same time scale, otherwise when *α*^2^ + *c* ≷ *r*, the prey reproduces slower/faster than the predator does.

**Figure S1:**
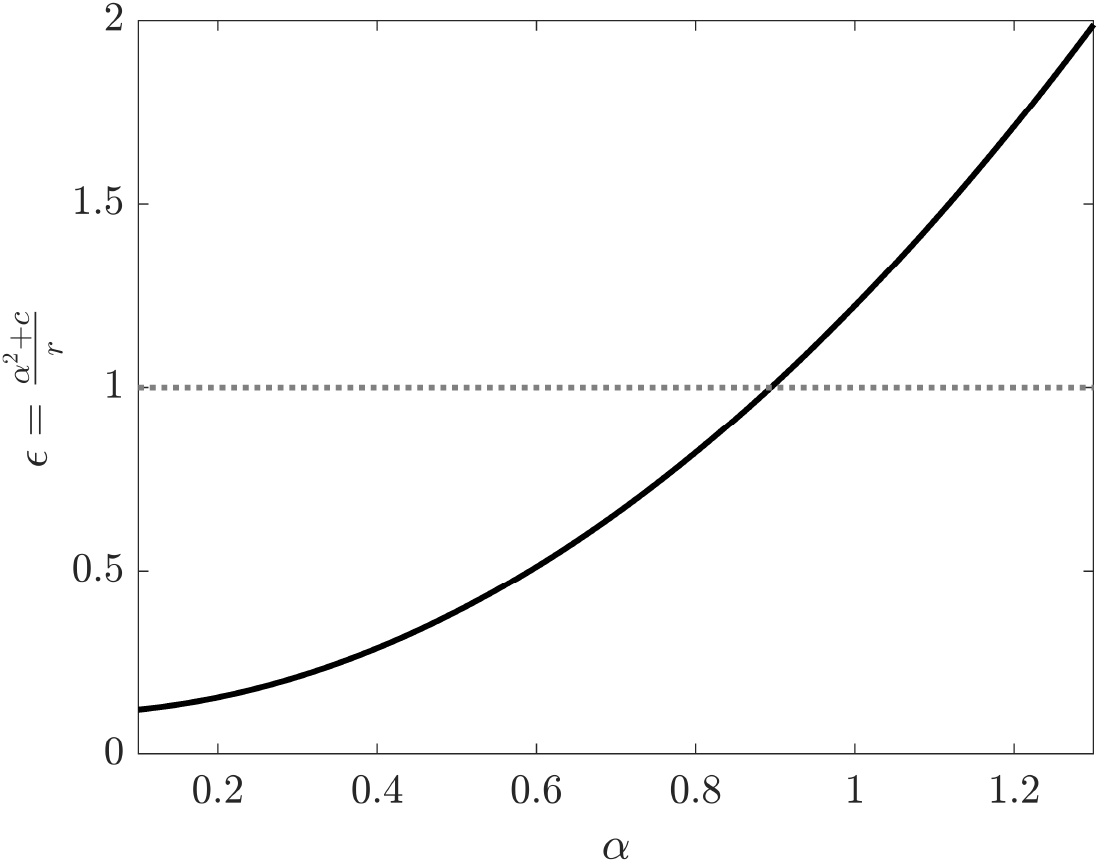
For mortality rate *m*(*α*) = *α*^2^ + *c*, the time-scale separation *ϵ* increases with increasing attack rate *α*. For *α* ≷ 0.9, the predator evolves faster/slower than its prey. For *α* ≈ 0.9, predator and prey evolve on the same time scale.

## S2 The Eco-evolutionary system

In this section, we study the stability of the long-term states of the eco-evolutionary predator-prey system (S7)–(S9):

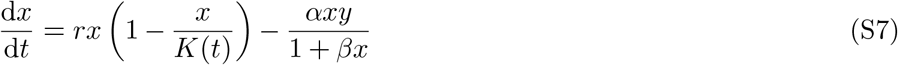

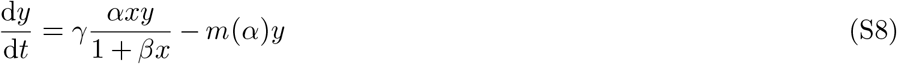

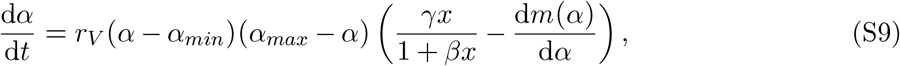

with time-dependent parameter *K*(*t*): *K*(*t*) = *max*(*K*_0_ − *r*_*K*_*t, K*_*min*_). Equation (S9) is zero if (i) *α* = *α*_*min*_, (ii) *α* = *α*_*max*_ or (iii) 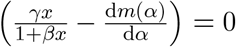 With *m*(*α*) = *α*^2^ + *c*, the eight equilibria ***e***_1_(*K*(*t*), *ϵ*) − ***e***_8_(*K*(*t*), *ϵ*) (S10)–(S17) are given by:

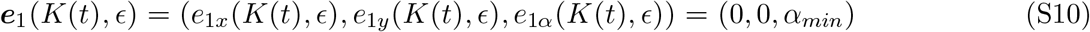

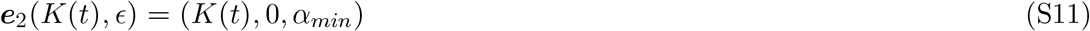

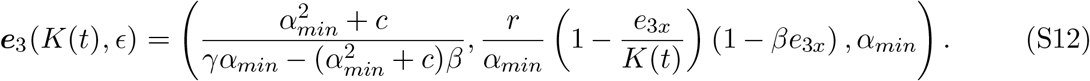

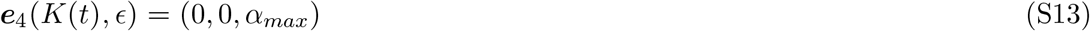

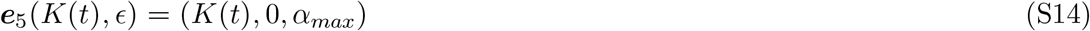

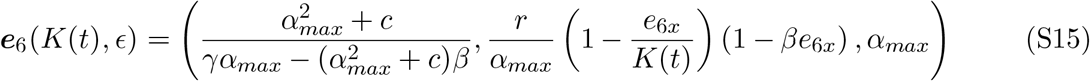

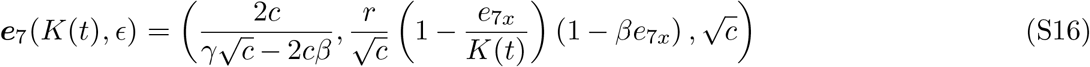

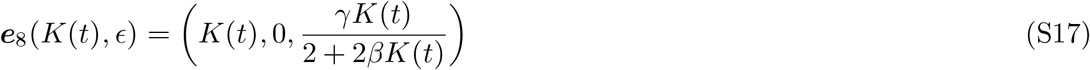

Figure S2 shows the stability of all eight equilibria ***e***_1_(*K*(*t*), *ϵ*) − ***e***_8_(*K*(*t*), *ϵ*) (S10)–(S17) depending on the time-dependent carrying capacity *K*(*t*) and the genetic variation *r*_*V*_. The equilibria ***e***_1_(*K*(*t*), *ϵ*)–***e***_6_(*K*(*t*), *ϵ*) are unstable ((*u*), orange) for all values of *K* and *r*_*V*_. Hence, the seventh ***e***_7_(*K*(*t*), *ϵ*) or eight equilibrium ***e***_8_(*K*(*t*), *ϵ*) represent the globally stable equilibrium of the system depending on *K*(*t*). For higher genetic variation *r*_*V*_, the stability range of the seventh equilibrium ***e***_7_(*K*(*t*), *ϵ*) extends even to higher values of *K*(*t*) (see fig. S2g).

**Figure S2:**
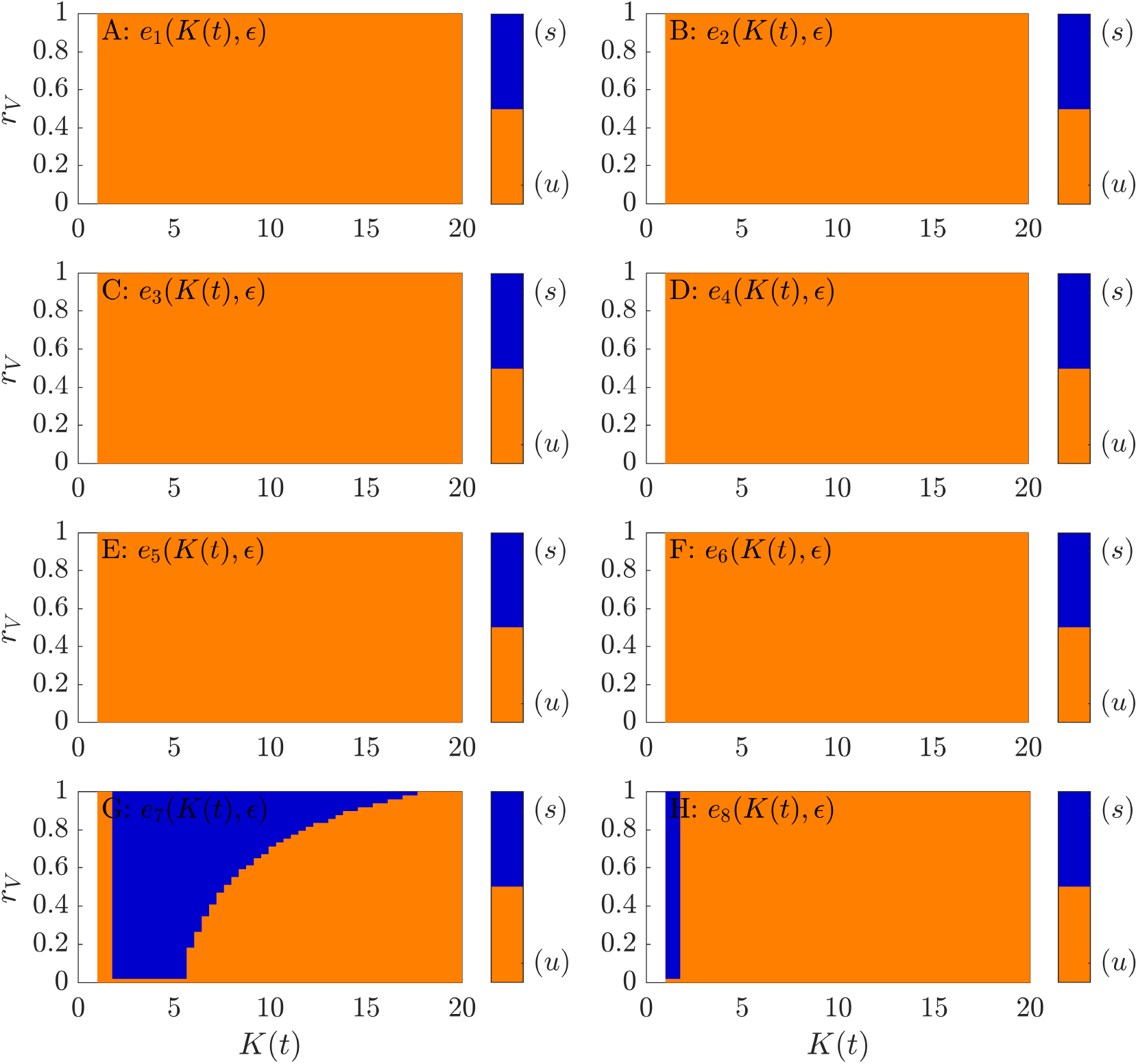
Stability of the eight equilibria ***e***_1_(*K*(*t*), *ϵ*)–***e***_8_(*K*(*t*), *ϵ*) (S10)–(S17) depending on the genetic variance *r*_*V*_ and the time-dependent carrying capacity *K*(*t*). Blue: stable (*s*), orange: unstable (*u*). Parameter: tab. 1

## S3 Eco-evolutionary dynamics for different trade-offs

In this section, we study if R-tipping and indirect evolutionary rescue still occurs for different trade-offs between predation 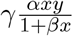 and the mortality of the predator *m*(*α*)*y*. Therefore, we introduce the trade-off parameter *k*: *m*(*α*) = *α*^*k*^ + *c* with *k* ∈ **R**_**0**_^+^. The eco-evolutionary system becomes:

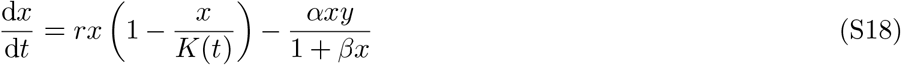

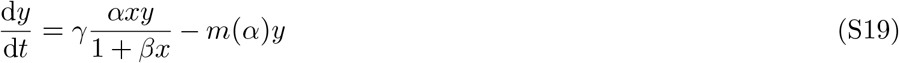

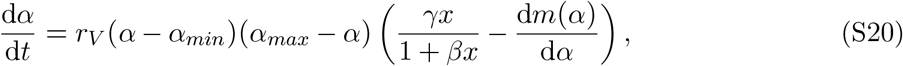

with time-dependent parameter *K*(*t*): *K*(*t*) = *max*(*K*_0_ − *r*_*K*_*t, K*_*min*_). As shown by Eq. (S19) and Eq. (S20), the trade-off parameter *k* affects directly (i) the mortality of the predator *m*(*α*) and especially, (ii) the fitness gradient 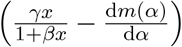. Hence, the parameter *k* influences the equilibrium (optimum) state ***e***_opt_(*K*(*t*), *ϵ*) of the eco-evolutionary system (S18)–(S20) for which the fitness gradient vanishes 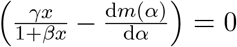.

Setting Eqs. (S18)–(S20) to zero and solving for the state variables *x, y* and *α* leads to the following eight equilibria ***e***_1_(*K*(*t*), *ϵ*)–***e***_8_(*K*(*t*), *ϵ*) (S21)–(S28):

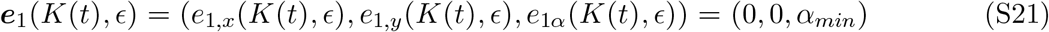

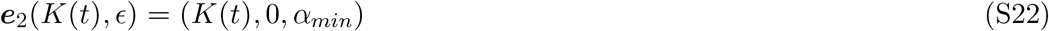

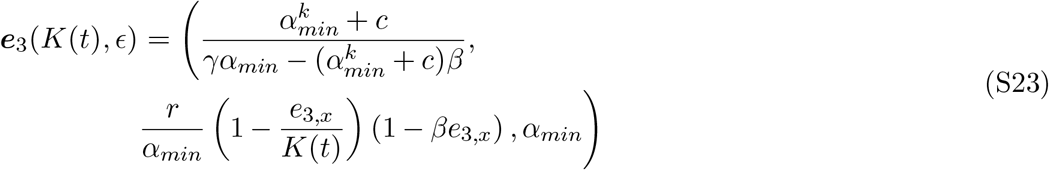

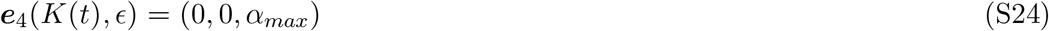

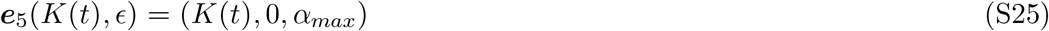

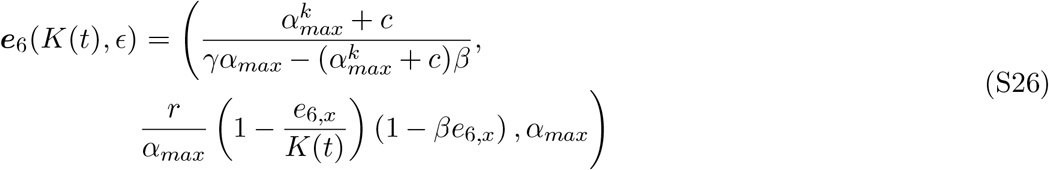

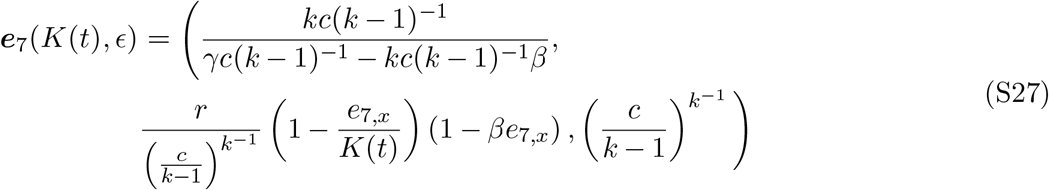

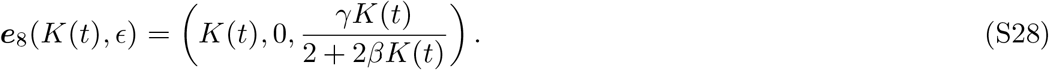

Notice, setting *k* = 2 leads to the optimum state ***e***_opt_(*K*(*t*), *ϵ*) (11) introduced in the section: *Indirect evolutionary rescue prevents rate-induced tipping*. There, the optimum state was represented by one of the two stable moving equilibria ***e***_7_(*K*(*t*), *ϵ*) and ***e***_8_(*K*(*t*), *ϵ*) depending on the value of *K*(*t*). Interestingly, for *k* ≥ 3, ***e***_7_(*K*(*t*), *ϵ*) (S27) represents the unique stable moving equilibrium for all values of the time-dependent carrying capacity *K*(*t*) because ***e***_8_(*K*(*t*), *ϵ*) (S28) is unstable (compare the colored regions above *k* ≥ 3 in fig. S3). Hence, the predator population is not extinct in the long-term ***e***_7_(*K*_*min*_, *ϵ*) (S27) (*e*_7,*y*_(*K*(*t*), *ϵ*) ≠ 0). Figure S4 demonstrates that we still find the rate-induced collapse of the prey and its prevention by indirect evolutionary rescue even though the predator persists in the long-term state ***e***_7_(*K*_*min*_, *ϵ*) (magenta dotted lines) for *k* = 3.

**Figure S3:**
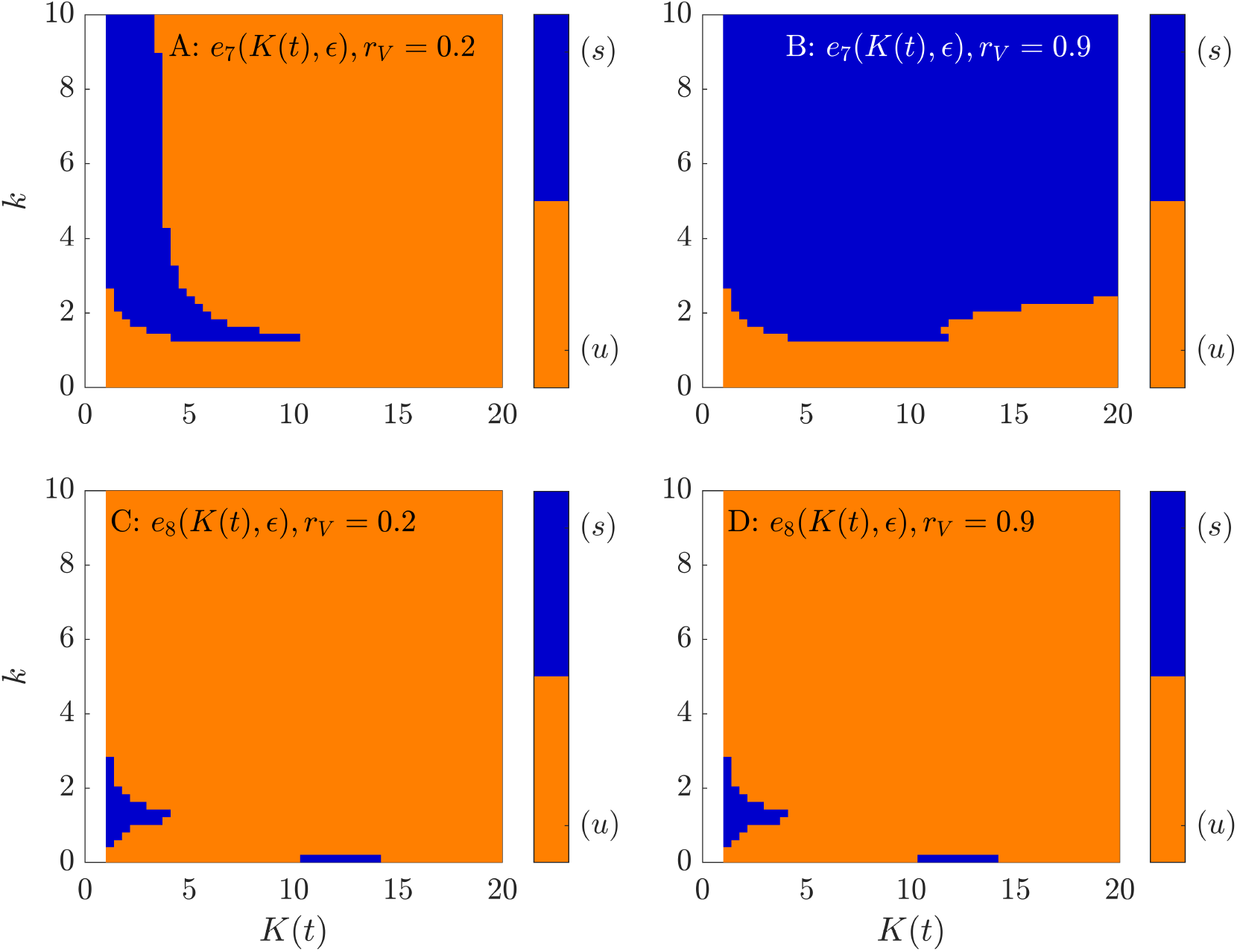
Stability of ***e***_1_(*K*(*t*), *ϵ*) (S27) and ***e***_8_(*K*(*t*), *ϵ*) (S28) depending on the trade-off parameter *k* and the time-dependent carrying capacity *K*(*t*). Blue: stable (*s*), orange: unstable (*u*). Parameters: tab. 1.

**Figure S4:**
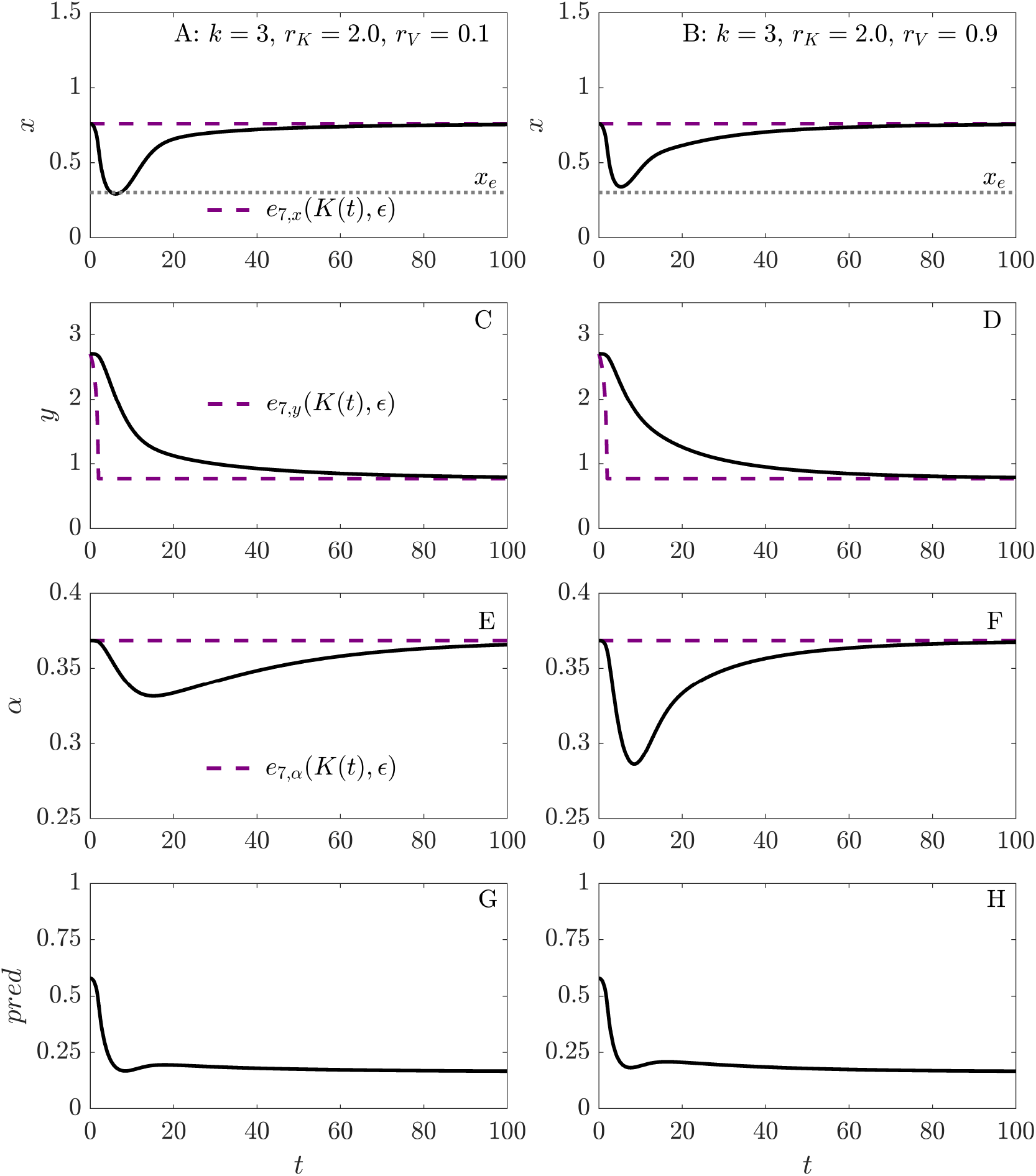
Prey density *x*, predator density *y*, attack rate *α* and predation pressure 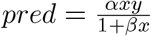 of the eco-evolutionary system (S18)–(S20) for *r*_*V*_ = 0.1 (left column) and *r*_*V*_ = 0.9 (right) column. The optimum state ***e***_7_(*K*(*t*), *ϵ*) (S27) is marked as magenta dashed line. Parameter: *x*_0_ = *n*_*x*_(*e*_7,*α*_),*n*_*y*_(*K*_0_ = 5.0,*e*_7,*α*_),*α*_0_ = *e*_7,*α*_(*K*(*t*),*ϵ*).Other parameters see tab. 1.

